# scVIVA: a probabilistic framework for representation of cells and their environments in spatial transcriptomics

**DOI:** 10.1101/2025.06.01.657182

**Authors:** Nathan Levy, Florian Ingelfinger, Artemii Bakulin, Giacomo Cinnirella, Pierre Boyeau, Boaz Nadler, Can Ergen, Nir Yosef

## Abstract

Spatial transcriptomics provides a significant advance over studies of dissociated cells in that it reveals the environment in which cells reside, thus opening the way for a more complete description of their state and function. However, most current methods for embedding and discovery of cell states rely only on the cells’ own gene expression profile, thus raising the need for ways to account for the neighboring cells as well. Here, we introduce scVIVA, a deep generative model that leverages both cell-intrinsic and neighboring gene expression profiles to output stochastic embeddings of cell states as well as normalized gene expression profiles. We demonstrate that scVIVA produces informative fine-grained partitions of cells that reflect both their internal state and the surrounding tissue and that its generative model facilitates the testing of hypotheses of differential expression between tissue niches. We leverage these properties of scVIVA to uncover a spatially-restricted tumor-promoting endothelial population in breast cancer and niche-associated T cell states that are shared across multiple cancers. scVIVA is available as open source software within scvi-tools.org.

## 1 Introduction

The function and molecular composition of cells in multi-cellular organisms are strongly influenced by their surrounding cellular environment. This phenomenon facilitates tissue-level functions, which rely on coordination between and within cellular compartments. A convenient way to describe the structure of tissues is through *niches* - spatially restricted areas that have a characteristic composition of cell types and states. Accordingly, cells of the same type may evoke different functions depending on the niches they populate, reflecting, for instance, different phases of development and activation (Yayon et al., 2024), different stresses (Buck et al., 2017), or different homeostatic functions (Yosef & Regev, 2016). Characterizing cells in the context of their close environment can therefore lead to a better understanding of the mechanisms of tissue organization and function (Wagner et al., 2016; Palla et al., 2022).

Emerging technologies for spatial transcriptomics at a single-cell resolution (ST) can measure the transcriptomes of individual cells while also recording their tissue coordinates, thus offering the possibility of interrogating the contribution of individual cells to higher-level tissue functions. Studying these questions, however, requires computational tools that describe the state of each cell while accounting both for its own transcriptome and its immediate environment. Such tools should have the capacity to derive an embedding of each cell that accounts for technical effects (e.g., batches, sampling depths) and reflects important intracellular features as well as properties of the cell’s immediate environment (henceforth, *neighborhood properties*). Another desirable capability is providing access to normalized (e.g., batch-corrected) gene expression profiles to allow for more accurate gene-level analysis (e.g., comparing niches).

Currently, most studies of ST data rely on methods for non-spatially resolved single-cell RNA-sequencing such as scVI (Lopez et al., 2018) or Harmony (Korsunsky et al., 2019), ignoring spatial relationships when defining cell populations (Cadinu et al., 2024). These methods learn a low-dimensional embedding space that is corrected for nuisance variables but accounts only for the cells’ own profiles. Recently, methods have been developed that learn a joint embedding of cellular state and spatial context. Haviv et al. (2024) introduced ENVI, a model that represents cells in a way that reflects the gene-covariance of their immediate neighborhood. While the model indeed learns a spatially aware embedding, it requires paired scRNA sequencing data, which can be challenging to obtain in many cases, e.g., due to issues with profiling certain subsets (owing to low numbers or to difficulties in dissociation and isolation). In contrast with ENVI, which is trained to *predict* the niche for the cellular state, other models including simVI (Dong et al., 2023) and BANKSY (Singhal et al., 2024), were built to *leverage* spatial information to learn the cell state while not requiring paired single cell profiles. Remaining caveats, however, are the ability to effectively integrate multiple samples, to correct bias and estimate noise in the observed expression levels, and to account only for the *relevant* neighborhood properties, namely ones that are associated with the cell’s transcriptome.

Here, we present scVIVA: a probabilistic deep learning tool for the analysis of ST data that accounts for the niche context and the expression profile of each cell. scVIVA (pronounce *ess-see-vee-vah*, from environment in Hebrew) design enables a broad set of analysis types, providing both low-dimensional embedding of cells as well as normalized gene expression values and a Bayesian strategy for testing hypotheses of differential expression. The method was designed to address the current major caveats. First, it relies only on ST data and facilitates accurate integration of different ST datasets. Second, it only accounts for niche properties that are associated with the cell’s transcriptome and therefore more relevant for defining cell state. We showcase scVIVA’s ability to detect subsets of cells that differ in their spatial niche by dissecting murine astrocytes into fine-grained regional subsets corresponding to astrocytes within manually annotated regions of the murine brain. In a second case study, we characterized a population of endothelial cells promoting angiogenesis in an invasive niche of human breast cancer. Finally, using scVIVA to analyze an atlas of 18 tumor slices from eight types of cancer, we uncovered a spatially-aware stratification of regulatory T helper cells into circulating and highly immunosuppressive phenotypes, each linked to different tumor microenvironments.

## 2 Results

### 2.1 Overview of scVIVA

scVIVA requires a single-cell resolved spatial transcriptomics dataset. For each cell *n*, we denote its gene expression vector as ***x***_*n*_, its spatial coordinates as ***y***_*n*_, and, optionally, its sample ID as ***s***_*n*_. We also leverage a user-provided (potentially coarse) cell-type label *c*_*n*_. scVIVA derives a representation of cells that capture their own transcriptome as well as a summary of the cellular and molecular composition of their environment. To this end, it defines the spatial context of each cell in two ways. The first is the cell-type composition of its cellular neighborhood, denoted ***α***_*n*_. The second is the average gene expression state of neighboring cells, with a separate profile for each of the present cell types, denoted ***η***_*n*_ (Figure 1A). These cell-intrinsic gene expression states are learned with a spatially unaware model, which is trained first. Throughout the manuscript, we use scANVI (Xu et al., 2021) as this first embedding model, and neighborhoods are defined as the *K* = 20 spatially nearest neighbors.

**Figure 1.**
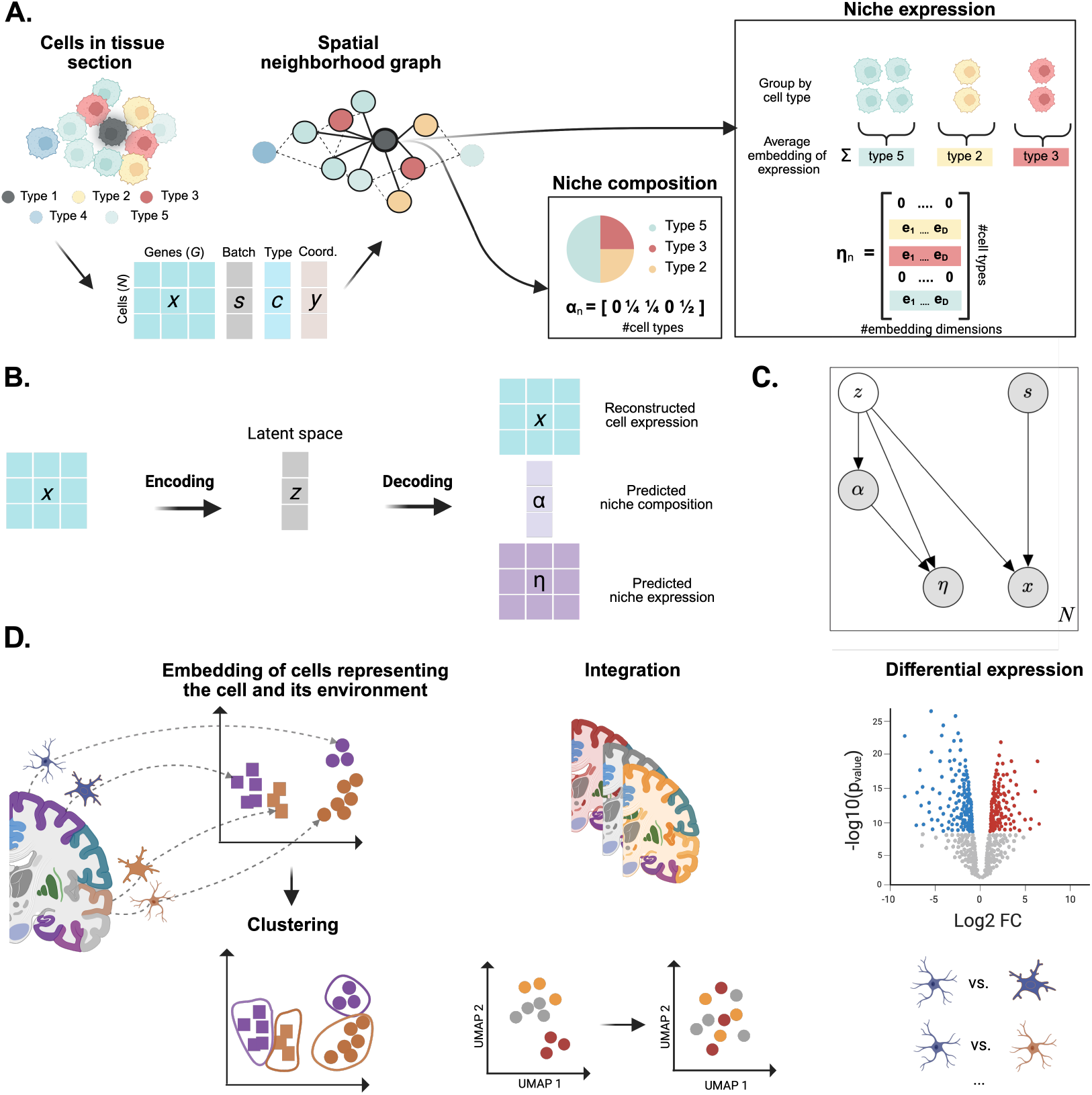
Overview of scVIVA. **A**. We are given a single cell resolution spatial transcriptomics dataset consisting of *N* cells and *G* genes, along with spatial coordinates and rely on prior cell type annotation and gene expression embedding using a spatially-unaware model. The cell niche is defined by the *K* nearest neighbors, from which the vector of cell type proportions is computed. We also retrieve the *K* gene expression embeddings of the neighbors and average them by cell type. If a cell type is not present in the niche, its entry is set to zero. **B**. scVIVA uses a probabilistic auto-encoder architecture to encode the cell’s observed gene expression, reconstruct the cell’s expression, and predict the composition and expression of its surrounding niche. **C**. The underlying graphical model. Shaded nodes represent observed random variables. Empty represent latent random variables. Edges signify conditional dependency. The rectangle represents independent replication across the *N* cells. **D**. Use cases of scVIVA: cell embeddings reflect the cell type and its environment. We can leverage these embeddings to cluster the cells by niche-dependent cell states. scVIVA corrects for batch effects and enables the integration of niche-dependent cell states across different tissue slides. The generative part of the model is used for differential expression analysis between cell types and cell states across niches.

scVIVA employs a latent variable model inferred with a variational auto-encoder (VAE). The trained model provides a probabilistic, low-dimensional representation of the state of each cell, ***z***_*n*_, that is corrected for batch effects and captures its gene expression profile and its neighborhood. From the low-dimensional embedding, the model predicts ***α***_*n*_ and ***η***_*n*_, and reconstructs 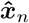. This design endows the embedding ***z***_*n*_ with knowledge about the surrounding of the cell (***α***_*n*_ and ***η***_*n*_), while at the same time limiting the influence of the tissue neighborhood only to properties that are predictable by the cell’s profile ***x***_*n*_. Moreover, the capacity to probabilistically generate normalized gene expression estimates 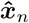 allows for testing hypotheses of gene expression changes (Figure 1B-C), while the embedding ***z***_*n*_ can be used to stratify cells into types and states, or along phenotypic gradients.

To set the loss functions used for training the model, the vector of gene expression of a cell, ***x*** is modeled with a Poisson distribution (Lopez et al., 2018), while the niche variables ***α*** and ***η*** are modeled with Dirichlet and Gaussian distributions, respectively (alternative implementations are readily available for users of our code base, e.g. negative binomial for ***x***). All three losses are optimized concurrently. The embedding variable ***z*** is modeled with a normal distribution, using a standard Gaussian as prior. The optimization procedure relies on four neural networks: an encoder, which estimates the cell state ***z*** from the observed counts ***x*** (estimating the variational posterior *q*(***z*** | ***x***)), an expression decoder that generates expression given a cellular state (estimating *p*(***x*** | ***z***)), a cell-type proportion decoder that estimates *p*(***α*** | ***z***), and a cell state decoder that estimates *p*(***η*** | ***z***). All the model parameters are learned via evidence lower bound (ELBO) maximization (Kingma & Welling, 2013).

The implementation of scVIVA was designed to utilize graphics processing units (GPUs), it can handle millions of cells and thousands of genes. The training time is similar to spatially-unaware models, with run times ranging from one hour for a medium-sized dataset (200k cells and 300 genes) to the order of 15 hours for a large dataset (10 million cells and 550 genes); using one NVIDIA RTX 4090 GPU; not including the time needed for calculating the spatially unaware embeddings. scVIVA is therefore capable of performing context-aware embedding of massive datasets. It allows spatially-aware analysis of cellular state, provides an integrated view across multiple samples, and provides capabilities for differential expression analysis between distinct niche-dependent cell states (Figure 1D). An in-depth description of scVIVA is provided in Methods.

### 2.2 SCVIVA DISTINGUISHES NICHE-DEPENDENT CELL STATES IN THE MOUSE BRAIN

As our first test case, we applied scVIVA to a dataset of ST samples consisting of a panel of 1100 genes across six coronal sections of the murine brain using MERFISH (two mice, each having three adjacent sections (Zhang et al., 2023)). In the following we treat each section as a separate batch. Our goal was to evaluate the ability of scVIVA to learn meaningful spatially aware embeddings of cells and compare it with previous approaches. We applied scVIVA and compared its latent space against scANVI (acting as a common non-spatial benchmark), simVI, BANKSY, and Nicheformer (either zero-shot or fine-tuned), using metrics from the *single cell integration benchmark* (scIB) (Luecken et al., 2022).

In our first analysis, we applied scIB in a manner decoupled from spatial information: we evaluated the extent to which cells of different batches are mixed (“Batch correction”) and at the same time, the extent to which cells from the same type are clustered together (“Cell type preservation”; using annotations provided by the authors of the dataset, Figure 2A). We found comparable performance across all methods, except Nicheformer and BANKSY, which had lower scores. Since BANKSY does not natively account for batch effects, we added to its pipeline a batch-correction component using Harmony (Korsunsky et al., 2019), as recommended by its authors. While it improved the score, the corrected embedding still yielded lower scIB scores compared to scVIVA.

**Figure 2.**
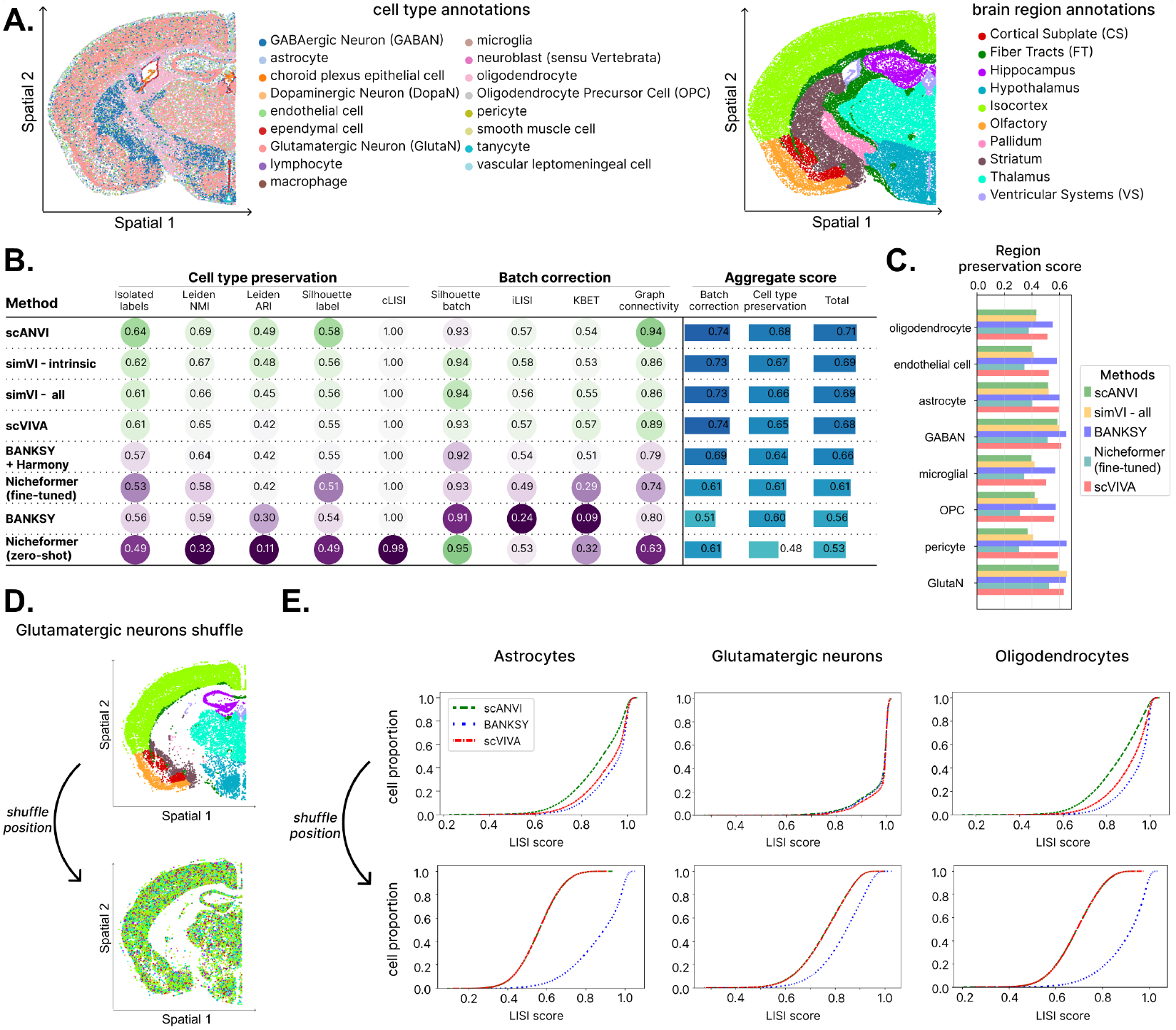
scVIVA learns a cell embedding reflecting niche-dependent cell states. **A**. Brain tissue slice colored by cell type labels (left) and by major regions (right). **B**. scIB metrics benchmarking scANVI, scVIVA, BANKSY, Nicheformer (zero-shot and fine-tuned), and different flavors of simVI: intrinsic embedding alone and concatenated with spatial-induced. The spatial-induced alone is not displayed as it is not meant to retain cell type information. In the following, the concatenated latent space is used for simVI as it performed best in those benchmarks. Likewise, we use fine-tuned Nicheformer. Raw metric scores are displayed. **C**. scIB metrics using the brain region annotation as label, when restricting the embedding to a single cell type. Only cell types with at least 400 occurrences in at least two regions are included, and the region preservation score (bio-conservation score in scib-metrics) is reported. **D**. We shuffle the cells spatial coordinates of all cells of a distinct cell type as exemplified for glutamatergic neurons. **E**. Cumulative distribution function of the LISI scores (Korsunsky et al., 2019) of each method before and after shuffling, for 3 example cell types (other cell-types in Supplementary Figure S1).

In our second analysis, we applied scIB in a manner that reflects the spatial context of cells. To this end, we considered each cell type separately and used the cells’ region as a substitute for the cell type label (using a stratification of the brain into its canonical regions, Figure 2A). This way, the “Cell type preservation” metrics of scIB capture the extent to which the embedding groups cells according to their surrounding tissue environment. Overall, we observed that BANKSY and scVIVA have the best performance in region label preservation, whereas simVI shows limited improvement over the non-spatial benchmark scANVI (Figure 2C). Notably, the limited performance of Nicheformer is likely due to the fact that spatial coordinates are not accounted for during training. Overall, this benchmark places scVIVA favorably compared to the other methods, striking a balance between integration and preservation of cell type and niche information.

scVIVA, similar to its benchmark methods, aims to learn a cell-centric representation; namely - a representation that reflects the state of the cell, while emphasizing properties that are impacted by the cell’s environment. Therefore, such a model should not be influenced by properties of the tissue environment that are unrelated to the cell’s state. To test this, we designed a semi-synthetic experiment in which we shuffle the spatial coordinates of cells (one cell type at a time), thus eliminating the dependence between gene expression and tissue environment (Figure 2D). We use this corrupted data as input for each of the models and retrieve cell embeddings. We compute the Local Inverse Simpson Index (LISI) (Korsunsky et al., 2019) to study whether the shuffled niche information is still represented in the embedding. For instance, the LISI measure quantifies the extent to which astrocytes that are randomly assigned to the same region tend to be embedded together (i.e. with LISI close to one). We then compare the LISI distributions for shuffled cells against the LISI scores before shuffling. Intuitively, spatially-aware embeddings are expected to yield higher scores before shuffling, and any method should collapse to a random distribution after shuffling. We compared scVIVA and BANKSY, considering scANVI as a non-spatial control. Before shuffling (Figure 2E-top), we observe higher LISI values for the embeddings of BANKSY and scVIVA, in line with Figure 2C results. After shuffling (Figure 2E-bottom), scVIVA and scANVI highlight that niche and cellular state are independent of each other, reflecting an essentially random distribution of LISI values. BANKSY, however, still tends to group cells according to their shuffled tissue position. This result is expected since BANKSY uses an a-priori concatenation of cell and niche features as input. It therefore does not necessarily detect niche-dependent differences in gene expression, but instead represents cumulative differences in gene expression and niche composition.

Overall, this case study demonstrates that scVIVA provides an effective tool for cell embedding that corrects for batch effects while reflecting variation due to both cell type (Figure 2B) and the surrounding tissue (Figure 2C). In doing so, scVIVA is capable of screening out properties of the surrounding tissue that are unrelated to the cell’s transcriptome, thus emphasizing niche-dependent properties of gene expression (Figure 2D-E).

### 2.3 SCVIVA DELINEATES REGION-SPECIFIC ASTROCYTE POPULATIONS IN THE MOUSE BRAIN

We next studied whether capturing niche information allows novel insights into cell states. We focused on astrocytes, a major type of glial cells present in different regions of the brain. These cells support neurons by maintaining the blood-brain barrier, facilitating neurotransmitter uptake, ionic balance, synapse formation, and synaptic transmission (Lee et al., 2022). There is growing evidence of regional specificity within astrocytes that is associated with distinct functional roles (Zhang et al., 2023; Yao et al., 2023; Endo et al., 2022). To investigate brain region-specific gene variation in astrocytes, we analyzed their spatially aware latent representations produced by scVIVA. In order to retrieve region-specific gene expression programs, we applied Hotspot (DeTomaso & Yosef, 2021) to identify modules of autocorrelated genes, where the distances between cells are defined by scVIVA embeddings (Methods). This analysis resulted in 14 modules (Figure S2), representing different gene expression programs in astocytes, most of which are highly region-specific (Figure S3), including modules that are largely restricted to the striatum, thalamus, and hypothalamus (Figure 3C). The thalamus module contained *Lgr6*, previously identified as a marker of murine medial thalamic astrocytes (Kleshchevnikov et al., 2022) as well as human thalamic astrocytes (Mathys et al., 2024; Kim et al., 2023a). In line with previous experiments (Herrero-Navarro et al., 2021), this localized module also contained *Tcf7l2*, a gene that is essential for astrocyte maturation and tends to be highly expressed in the thalamus (Szewczyk et al., 2024). The hypothalamus module is characterized by *Igsf1* and *Itih3*, both known to be markers of astrocytes in the median eminence of the hypothalamus (Joustra et al., 2015; Pfau et al., 2024). The striatal module contained dopamine receptor D2 (*Drd2*,) which is indeed expressed by striatal astrocytes (Amato et al., 2023); and *µ*-crystallin (*Crym*), a known marker of striatal astrocytes (Kim et al., 2023b; Ollivier et al., 2024; Khakh, 2019).

**Figure 3.**
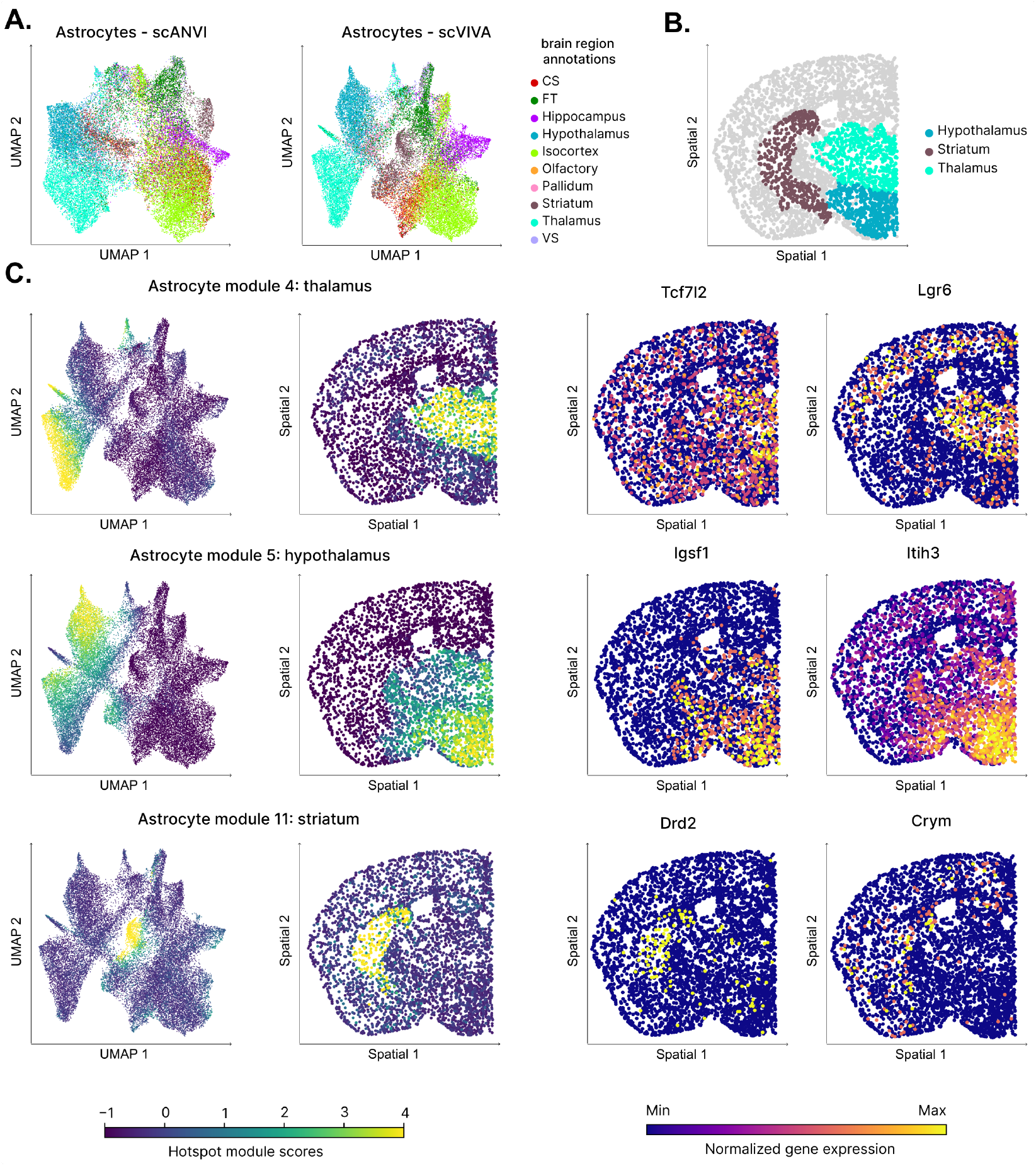
scVIVA stratifies astrocytes into niche-dependent cell states. **A**. Comparison of scVIVA (right) and scANVI (left) based UMAPs for astrocytes colored by region labels. **B**. Spatial plot of astrocytes colored by region labels. **C**. Using Hotspot (DeTomaso & Yosef, 2021), each region can be associated with a gene module. Module scores, representing the activation of the module genes in a cell, are displayed in UMAP space and spatial coordinates (left). For each module, we showcase region-specific astrocyte markers (right).

To compare scVIVA’s ability to retrieve spatially-organized gene programs, we used Hotspot to generate gene modules from the spatially-unaware embeddings of scANVI (Figure 3A-left, Figure S4). We manually aligned modules identified with both methods and computed the spatial autocorrelation of the module scores. We found significantly greater values for scVIVA modules (Wilcoxon signed-rank test, *p <* 0.05). In addition, scVIVA shows significantly higher ROC-AUC values when using module scores to predict the main region labels (Wilcoxon signed-rank test, *p <* 0.05, Methods).

We conclude that the spatially aware scVIVA model enables the stratification of cells not only by their type but also by their niche content. Using astrocytes as an example, we identified cell-type specific and spatially restricted gene expression programs, highlighting different brain regions and recovering well-studied and recently identified markers.

### 2.4 Differential expression testing with scVIVA REVEALS NEO-VASCULARIZATION IN TUMOR REGIONS OF BREAST CANCER

In addition to informative low-dimensional embedding, scVIVA provides normalized estimates of gene expression values as well as tests for differential expression (DE) via its generative model. To demonstrate these utilities, we applied scVIVA to an in-situ sequencing dataset of a human breast cancer section, generated with 10X Xenium (Janesick et al., 2023). This dataset comprises 313 genes and 280k cells, covering various immune, stromal and malignant subsets (Figure 4A). We found that the cell segmentation originally performed includes many erroneously assigned transcripts and we therefore re-segmented the data using Proseg (Jones et al., 2024), leading to improved cell-type delineation (Figure 4B). In the following, unless specified, we will report results from the re-segmented data.

**Figure 4.**
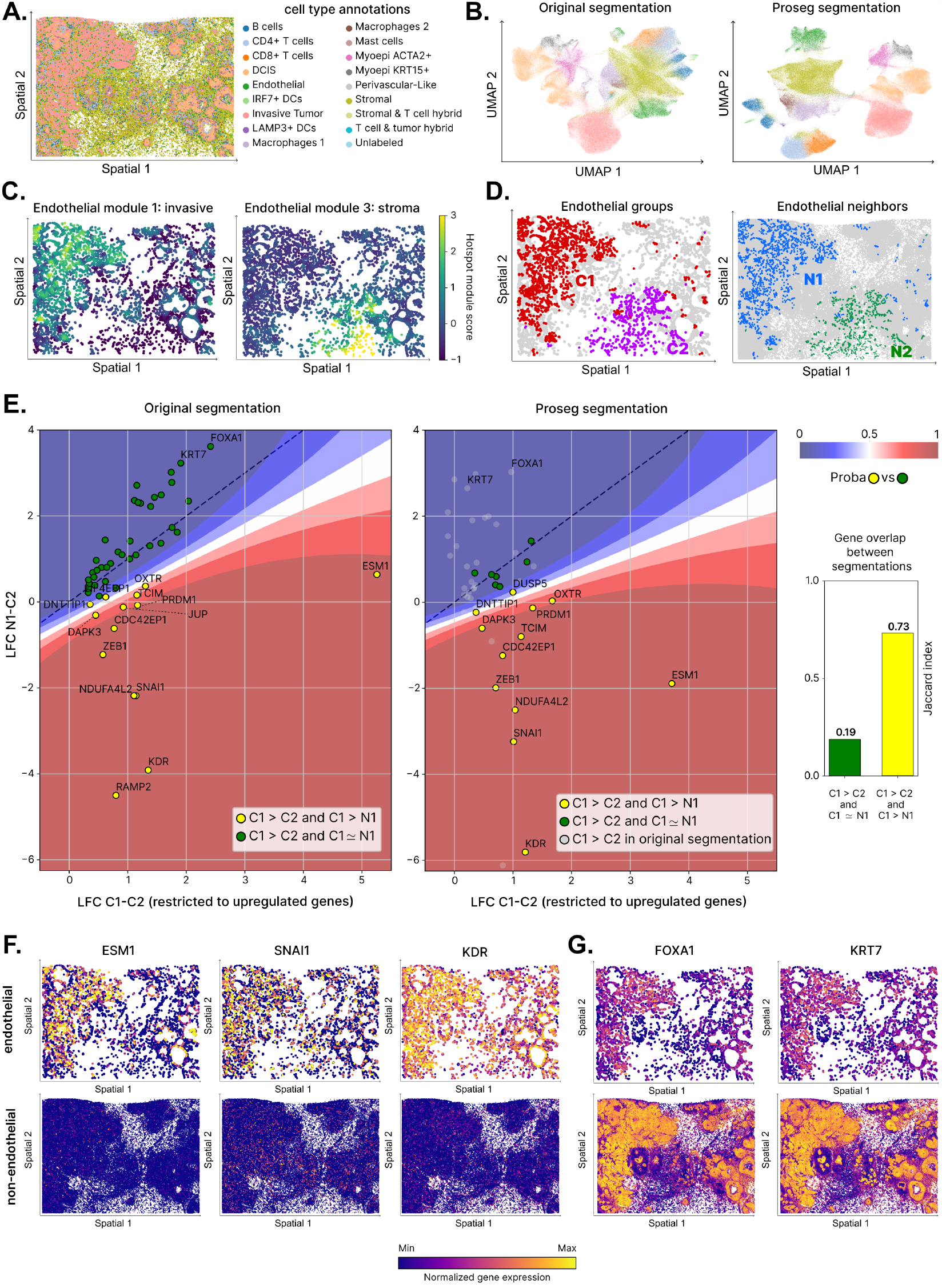
scVIVA enables the characterization of spatially confined endothelial populations in breast cancer samples. **A**. Tissue slice colored by cell type labels. **B**. UMAP of scVIVA embeddings, using the original cell segmentation (left) and after re-segmenting the cells using Proseg (right). Cells are colored by cell type labels. **C**. On the re-segmented data, endothelial gene modules from Hotspot applied on spatial coordinates. One module co-localizes with the invasive tumor region and the other one with the tumor stroma. Cells are colored by module scores. **D**. We identify endothelial cells in the invasive niche (*C1*) and endothelial cells in the stroma (*C2*) (Methods). *N1, N2* refers to the nearest spatial neighbors of *C1* and *C2* while ignoring endothelial cells. **E**. Median Log-Fold Change (LFC) of upregulated genes in *C1* vs *C2*, using the original segmentation (left) and the re-segmented data (right) displayed on the x-axis, while we compare differential expression computed between *N1* and *C2* on the y-axis. Genes are colored by their marker label. The classifier decision boundary is shown, and the dashed line represents the identity line. Gene overlaps for the sets of markers/ neighborhood genes are measured by Jaccard index. **F**. Spatial gene expression of marker genes upregulated in both segmentations, subset to endothelial cells (top) and subset to non-endothelial cells (bottom). These genes are higher expressed in endothelial cells. We display the observed expression based on the Proseg segmentation; corresponding spatial plots using the original segmentation are shown in Figure S6A. **G**. Similar to F. These genes are higher expressed in non-endothelial cells. Spatial plots from the original segmentation are shown in Figure S6B.

The authors of this dataset identified distinct regions of this specimen, corresponding to in situ ductal carcinoma (DCIS), invasive tumor and stroma. In DCIS, tumor cells are retained within the ductal–lobular system of the mammary gland, while in invasive tumors, the neoplastic cells penetrate the adjacent tissue (Casasent et al., 2022). Stromal regions are defined as the surrounding parts outside the main tumor regions. Here, we used scVIVA to study how cells of the same type act differently in the stromal and invasive environments. We focused our analysis on endothelial cells, which play crucial roles in the tumor microenvironment (TME). In some settings, these cells can help facilitate anti-tumor immunity by recruiting effector T and B cells (Harris et al., 2024). In other settings, these cells can help establish a tumor-supportive environment, e.g., through altered expression of cell adhesion and cell migration molecules or secretion of angiogenic factors (Sobierajska et al., 2020; Harris et al., 2024).

The partition of the tumor into distinct regions, reflecting stromal or invasive characteristics, is readily observable by the local composition of cell types (Figure 4A), and also in a Hotspot analysis of spatially autocorrelated gene modules (Figure S5). To identify the gene expression programs that may drive region-specific function of endothelial cells, we leveraged the generative capacity of scVIVA for differential expression (DE) testing (endothelial cells in invasive vs. stromal regions; *C1* and *C2* respectively in Figure 4D-left). Our DE procedure builds on lvm-DE, a general strategy we previously developed for comparative analysis with latent variable models (Boyeau et al. (2023); see Methods). Despite the re-segmentation we find persistent issues of molecule misassignment due to remaining errors in segmentation, which may lead to spurious DE results if we analyze the data as-is. We therefore developed a more refined DE procedure to mitigate this limitation. We defined two additional sets of cells: *N1* and *N2*, the non-endothelial spatial nearest neighbors of *C1* and *C2*, respectively (Figure 4D-right). We then used lvm-DE to compare *C1 vs. N1, N1 vs. C2, N2 vs. C2* and *N2 vs. C1*, to identify genes whose DE signal can be explained by the endothelial environment. When looking at the upregulated genes of *C1*, we hypothesized that if a gene that has higher expression in *C1* vs. *C2* also has higher expression in *N1* vs. *C2*, then we cannot exclude that its signal in the former test is spurious. To distill posterior estimates of a gene being true DE based on the two tests (i.e. population of interest: *C1 vs. C2* and its environment: *N1 vs. C2*) we used a Gaussian process classifier (Figure 4D-left). The classifier is trained to identify genes that are clear endothelial markers (Methods).

We first investigated genes enriched within endothelial cells of the invasive region compared to the tumor stroma. To evaluate our DE procedure, we compared the results obtained with Proseg to those obtained with the original (less accurate) segmentation (Figure 4E). Generally, we expect that with a conservative method both segmentations will lead to similar results. Conversely, we expect that a simpler DE method will be more dependent on the segmentation quality, with the less accurate (with more transcript misassignment) segmentation yielding spurious results. Inspection of individual genes that are declared significant further supports the merit of the two-step DE approach compared to the simple (*C1* vs. *C2* only) approach. For instance, the simple approach identifies *FOXA1* (Balsalobre & Drouin, 2022; *Janesick et al*., *2023; Bhat-Nakshatri et al*., *2024)) and KRT7* (Elmentaite et al., 2022) that are normally expressed by epithelial cells to be enriched in the endothelial compartment of the invasive region. Indeed, spatial plots of these two genes show high expression in non-endothelial cells (Figures 4G, S6B). Conversely, the 2-step approach (using either segmentation method) identifies more plausible gene expression changes that are characteristic to the invasive tumor. Specifically, in both segmentations, the 2-step DE identified increased expression of ESM1, SNAI1, and *KDR*. ESM1 is an endothelial marker previously found to be upregulated in invasive breast cancer (Zeng et al., 2023). *SNAI1* is an endothelial-mesenchymal transition (EndMT) transcription factor and a key regulator of dysfunctional blood vessels in cancer (Hoffmann et al., 2024; Cabrerizo-Granados et al., 2021). Upon activation during EndMT, SNAI1 induces the expression of multiple other transcription factors, resulting in neoangiogenesis and tumor growth (Youssef & Nieto, 2024). The *KDR* gene encodes the vascular endothelial growth factor receptor 2 (VEGFR2), which is essential for angiogenesis and increased permeability (Pérez-Gutiérrez & Ferrara, 2023). Spatial plots of *ESM1, SNAI1* and *KDR* show increased and more localized expression in endothelial cells (Figure 4F-top) compared to non-endothelial (Figure 4F-bottom). Taken together, gene expression changes highlighted by scVIVA and the 2-step DE indicate dysregulation of angiogenesis that is spatially confined to the invasive tumor region (Motzer et al., 2020; Li et al., 2022).

In contrast, endothelial cells located in the stroma express higher levels of canonical markers of endothelial cells (Figure S6C). For instance the 2-step DE identified *EDN1*, which is critical for vasoconstriction (Geldhof et al., 2022)), *CAVIN2* a regulator of caveolae and nitric oxide production that also influences vasodilatation (Aitken et al., 2023; Boopathy et al., 2017) and the canonical marker of endothelial cells *CLDN5* (Reed et al., 2024) (Figure S6D-E). Higher expression of these endothelial markers highlights more mature functional endothelial cells in the stroma compared to those in the invasive cancer region, which instead exhibits properties of endothelial-to-mesenchymal transition, including loss of markers of differentiated endothelium.

### 2.5 SCVIVA INTEGRATES T CELL NICHES ACROSS CANCER TYPES

To demonstrate the ability of scVIVA to extract insights from large (multi-sample) spatial transcriptomics datasets, we analyzed a MERFISH dataset from Vizgen encompassing 18 cancer tissue samples from eight different cancer tissues (Figure5A-left). This dataset comprised 550 genes and 10 million cells, which we annotated into coarse cell types (Figure 5A-right, Methods). Spatial visualization of cell type labels shows an expected organization into tumor-rich regions surrounded by immune-rich stroma regions (Figure 5B).

**Figure 5.**
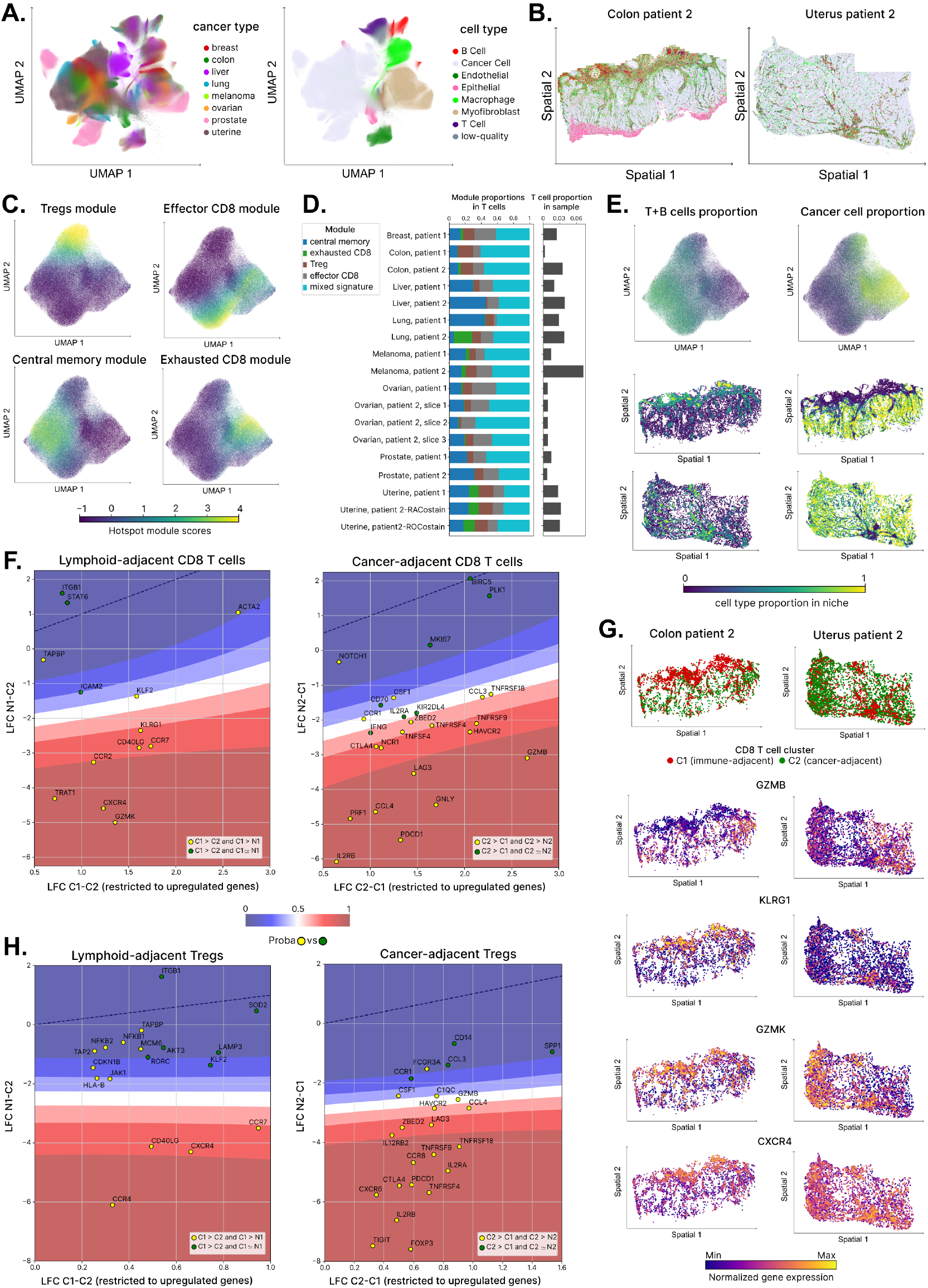
scVIVA integrates niche-dependent cell states across cancer types and patients. **A**. UMAP of scVIVA embeddings, colored by cancer type and cell type. **B**. Example tissue slices colored by cell type. **C**. UMAP of scVIVA embeddings restricted to T cells, colored by gene module scores computed using Hotspot on the scVIVA latent space. Four major T cell subtype-associated modules were identified, while other modules exhibited mixed phenotypes. **D**. T cell composition across patients. Each T cell is assigned to the module with the highest score unless the maximum score is below 0.5 or corresponds to a mixed phenotype module, in which case it is labeled ‘mixed signature’. The fraction of T cells within each sample is also displayed, highlighting substantial variability across patients. **E**. UMAP of T cell embeddings colored by niche composition fractions, specifically (***α***_T cell_ +***α***_B cell_) and ***α***_Cancer cell_ (top). Using the same color scheme, spatial distributions of T cells in two representative tissue samples are shown to highlight immune-adjacent and cancer-adjacent niches (bottom) **F**. Differential expression between two groups of cytotoxic T cells, obtained by k-means clustering on scVIVA latent (k=2). We identify immune-adjacent cytotoxic T cells as *C1* and cancer-adjacent as *C2* (Methods). *N1, N2* refers to the nearest neighbors in space of *C1* and *C2* while ignoring T cells. Left: median Log-Fold Change (LFC) of upregulated genes in *C1* vs *C2*, displayed on the x-axis, while we compare differential expression computed between *N1* and *C2* on the y-axis. Genes are colored by their marker label. The classifier decision boundary is shown, and the dashed line represents the identity line. Right: The same analysis as on the left, but comparing *C2* vs. *C1*. **G**. In two representative samples, spatial plots of *C1* and *C2* and spatial gene expression of selected markers. **H**. Similar analysis as in panel F, but for regulatory T cells (Tregs). Immune-adjacent Tregs are labeled as *C1*, and cancer-adjacent Tregs as *C2*.

To showcase scVIVA as a tool for integrated analysis of these tissues, we focused on the T cell compartment. First, we note that the overall numbers of T cells varied substantially between types of cancers and between patients of the same cancer type. For example, we observe a vastly different abundance of T cells between the two melanoma patients and very few T cells in ovarian cancer (Figure 5D), the latter reflecting the distinction between immune-hot and -excluded tumors (Galon & Bruni, 2019; Wu et al., 2024). We next applied Hotspot on scVIVA latent space in order to identify distinct and spatially organized gene expression programs in T cells. Four of the programs corresponded to readily interpretable T cell subsets: a regulatory CD4+ T cell program *(FOXP3, CD25, CTLA4*), a central memory T cell program (*CCR7, CD28, KLF2*), an effector CD8+ T cell program (*NKG7, GZMH, GZMK*), and an activation/exhaustion CD8+ T cell program (*PDCD1, LAG3, HAVCR2*; Figures 5C and S7C). Further supporting the variability between samples, we observe different frequencies of these subsets, even within the same type of cancer, e.g., in the proportions of exhausted CD8+ T cells in the two lung cancer patients (Figure 5D).

Considering the immediate environment of each cell (encoded by the observed niche composition ***α***; Figure 1A), the latent space of scVIVA clearly divided the T cell population into two groups: lymphoid-adjacent and tumor-adjacent (Figure 5E). Interestingly, the Treg and effector CD8+ T cell populations were present in both intratumoral and extratumoral (lymphoid-rich) niche types (Figure 5C-E). Conversely, the exhausted CD8+ T cell population was largely restricted to the tumor-rich niches across all tumor types investigated, thereby highlighting the association between immune infiltration and decrease of anti-tumor efficacy in cytotoxic T cells as a mechanism of tumor immune escape (Kersten et al., 2022). To explore the effect of the immediate environment on the function of Treg and CD8+ T cells (both effector and exhausted), we used the 2-step DE function of scVIVA. We compared, separately for the two subsets, cells in tumor-rich vs. lymphoid-rich environments.

Tumor-adjacent CD8+ T cells were enriched for immune checkpoint and exhaustion markers, including *HAVCR2* (encoding TIM-3), *LAG3*, and *PDCD1* (encoding PD-1), as well as cytotoxicity-related transcripts such as *GZMB* and *GNLY* (Figure 5F-left). Notably, tumor-adjacent CD8+ T cells also showed enrichment for *ZBED2*, a transcription factor recently implicated in the regulation of cytotoxicity and exhaustion-associated genes (Cillo et al., 2024), as well as *CCL3*, a chemokine expressed by both exhausted and effector memory CD8+ T cells (Buggert et al., 2023). In contrast, immune-adjacent CD8+ T cells were enriched for migratory chemokines receptors such as *CXCR4* and *CCR7*, and expressed cytotoxicity-related transcripts such as *GZMK* and *KLRG1* (Figure 5F-right). The expression of *CXCR4* has been shown to facilitate the retention of effector and memory CD8+ T cells at the tumor periphery and promote their egress via interactions with CXCL12-expressing peritumoral lymphatic vessels (Steele et al., 2023). Similarly, *CCR7* has been demonstrated to be involved in tissue egress and remigration to lymphoid organs (Bromley et al., 2005). Overall, integrated analysis of different cancers with scVIVA helped uncover spatially-associated segregation of the CD8+ population that is consistent across multiple tumor types (Figure 5G).

Similarly as in the cytotoxic T cell compartment, we observed that the lymphoid-adjacent Tregs displayed upregulation of migratory receptors *CCR7* and *CXCR4*, involved in lymphatic vessel-mediated egress (Figure 5H-left). In contrast, cancer-adjacent Tregs showed a clear tumor-infiltrating signature. Alongside the upregulation of inhibitory immune checkpoints such as *CTLA4, HAVCR2* and *PDCD1*, which were also enriched in the cancer-adjacent CD8+ T cells, we found overexpression of *FOXP3*, the master regulator of the immunosuppression gene program in Tregs (Li et al., 2020). We also detected enrichment of the co-inhibitory molecule *TIGIT*, which has been suggested to suppress pro-inflammatory Th1 and Th17 helper cells (Joller et al., 2014). Furthermore, tumor-infiltrating Tregs showed enrichment of the chemokine receptor *CCR8*, which emerged as a promising therapeutic target due to its specificity to tumor-infiltrating Tregs (Chen et al., 2024; Wen et al., 2025) (Figure 5H-right). Thus, spatial segregation of Tregs phenotypes across multiple cancer types reveals functionally distinct subsets: tumor-adjacent Tregs cells exhibit a highly immunosuppressive profile, while lymphoid-adjacent Tregs cells display characteristics associated with recirculating potential.

Overall, we identified niche-associated functional differences in T cells consistent across multiple cancer types. For both CD8+ and regulatory T cells, scVIVA successfully captured the stratification of lymphocytes into either an exhausted/immunosuppressive phenotype or a central memory-like phenotype, reflecting adaptations to their distinct microenvironments.

## 3 Discussion

Spatial transcriptomics provides a way to explore how the tissue environment affects gene expression and to more finely delineate different cell states. There is therefore a need to develop ways to define cell states in a manner aware of the spatial context and investigate state-specific spatially constrained gene expression programs while accounting for measurement noise and batch effect. Here, we develop scVIVA, a probabilistic algorithm that infers a stochastic latent representation from cells’ gene expression. From this latent space, we generate estimates of cells’ gene expression probabilities and predict a summary of each cell’s environment. We use a condensed representation of the environment, consisting of cell-type composition and average cell-type specific quantitative state, accounting for all cells within a fixed neighborhood. We demonstrate that adding the niche composition and niche state discriminative tasks into our non-spatial VAE framework improves the detection of niche-dependent cell states.

scVIVA can be used in diverse biological contexts and scales - ranging from a few replicates of the same tissue to massive multi-sample multi-tissue datasets. It provides access to fundamental functions. First, it produces cell embeddings that reflect both intrinsic gene expression and local niche properties that are statistically associated with (or rather, predictable by) the cell’s transcriptome. In the mouse brain, for example, scVIVA distinguishes astrocyte states across distinct regions such as the thalamus, hypothalamus, and striatum, revealing region-specific phenotypes. Second, scVIVA enables integration across multiple samples or tissues. By incorporating niche properties, we can identify shared and distinct niche-dependant cell states across samples. For instance, scVIVA uncovered a pan-cancer spatial heterogeneity of Tregs, highlighting a gradient in the immunosuppressive program from the tumor periphery to the core. Discovering such spatially regulated phenotypes can help uncover mechanisms by which cancer cells suppress anti-tumor immunity. Finally, the generative part of scVIVA enables differential expression testing. We introduced a principled method to filter out spurious signals when testing for differences between tissue regions and demonstrated that this approach consistently improves results and reduces spuriously detected genes. For instance, testing for differential expression between endothelial cells in the invasive and stromal niches of human breast cancer, our procedure identified the signature of immature endothelial cells with markers of angiogenesis while filtering out genes expressed by nearby epithelial cells.

The ongoing and anticipated assembly of large-scale spatial atlases with concomitant rich clinical metadata will help enable clinical discovery, for instance by associating patient-specific niche structure with diagnostic and prognostic covariates. We expect scVIVA to make an effective solution for driving such discoveries. scVIVA is integrated into the scvi-tools framework for open-source software and its broader scverse ecosystem, allowing extension with common downstream tasks like transfer mapping (Xu et al., 2021). scVIVA enables more robust and sensitive discoveries in spatial transcriptomics and we envision it to interact with sensitive tools for comparative analysis like multiple-instance learning (Litinetskaya et al., 2024) and differential abundance analysis (Boyeau et al., 2024), putting it at the forefront of discovering spatially regulated cell states.

## Methods

### A SCVIVA DETAILS

#### Single-cell spatial transcriptomics data

Consider a single-cell resolved spatial transcriptomics dataset, which provides the following measurements for *N* cells over *G* genes. For each cell *n*, we denote by ***x***_*n*_ *∈* ℕ^*G*^ the vector of observed gene expression counts in the cell. We assume that each cell is annotated, and we denote by *c*_*n*_ its discrete type assignment, taking values in 𝒯 = {1, …, *T*}, where *T* is the total number of cell types in the dataset. For each cell *n*, we also assume knowledge of batch annotations (donor, sample ID, etc.) stored in a vector ***s***_*n*_.

In addition, the experiment provides cell coordinates for each cell, ***y***_*n*_ *∈* ℝ^2^. We take the *K* nearest neighbors of a cell to define its niche using the Euclidean distance in physical space. We characterize the niche by its cell-type composition and gene expression. We denote by ***α***_*n*_ the *T* dimensional vector of cell type proportions among the *K* nearest neighbors of the cell *n*. Its values are in the probability simplex. For the surrounding cells in the niche, rather than using their full *G*-dimensional gene expression, we leverage a low dimensional gene expression embedding (PCA, scVI or similar) to represent each cell by a *D*-dimensional vector (*D ≪ G*). The niche gene expression is then defined as the average embedding expression of each cell type present in the niche. This is stored in a matrix ***η***_*n*_ *∈* ℝ^*T ×D*^ (Figure 1A).

#### Descriptive latent variable model

We propose a latent variable model for spatial transcriptomics data, aiming to capture both gene expression heterogeneity and spatial variation resulting from the micro-environment. In particular, we assume these two sources of variability are both captured by a *P* -dimensional latent variable (*P ≪ G*):

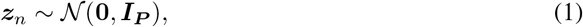

As our model is assumed to hold for any cell *n*, to simplify notations in what follows we drop the index such that {***z***_*n*_, ***s***_*n*_, ***α***_*n*_, ***x***_*n*_, ***η***_*n*_} = {***z, s, α, x, η***}. The random vector ***z*** captures both the cell expression and the *compositional* and *expression* variability of neighboring cells. We model the observed expression profile ***x*** as follows:

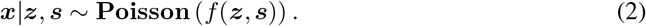

Here, *f* is a non-linear function with output in (ℝ^+^)^*G*^. We assume *f* is a composition of Multi-Layer Perceptrons (MLP), as in scVI (Lopez et al., 2018).

The latent variable ***z*** also captures the niche composition. We assume that the cell-type proportions of the cell’s *K* nearest neighbors are distributed as

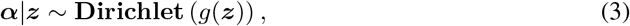

where *g* is a non-linear function with output in (ℝ^+^)^*T*^. We also parameterize *g* with a MLP. Last, we assume that the neighboring cells’ average expression profiles are distributed as

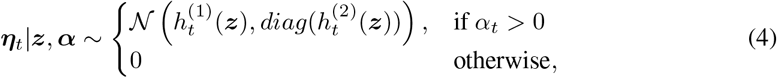

where *t* = 1, …, *T*. Therefore, the random variables 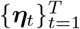 are conditionally independent given ***z, α***. The non-linear functions *h*^(1)^, *h*^(2)^ have outputs in ℝ^*D*^ and (ℝ^+^)^*D×D*^ respectively.

Given the observed data 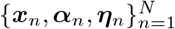 and their batch annotation 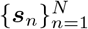, our goal is to learn the functions *f, g, h*^(1)^, *h*^(2)^ of the latent variable model above (Eq. 2-4). To this end, we adapt a variational autoencoder (VAE) framework for gene expression data to account for the niche observations. As part of this model, we also learn an encoder that estimates the cell state ***z*** from the observed counts ***x***. We rely on variational inference to derive a lower bound of the evidence log *p* (***α, x, η*** | ***s***).

#### Variational inference recipe

The evidence of the data can be decomposed as:

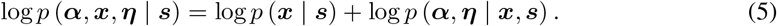

The first term in Eq. 5 corresponds to a generative model of the cell expression ***x***. The second term is a discriminative model of the niche properties (***α, η***) given the cell expression.

The lower bound for log *p* (***x*** | ***s***) is the standard evidence lower bound (ELBO) of a conditional VAE, with observed data ***x***|***s***. Letting *q* (***z*** | ***x, s***) denote the variational approximation of the posterior *p*(***z***|***x, s***), the ELBO is:

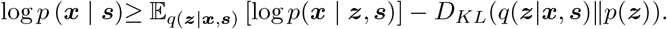

Next, we compute the lower bound for the niche term in Eq. 5. By the law of total probability:

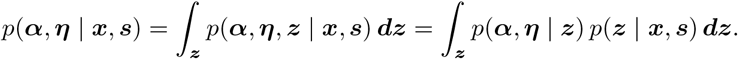

Replacing by the variational approximation, we get:

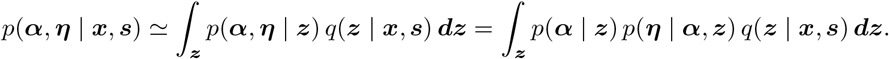

Hence, by Jensen’s inequality,

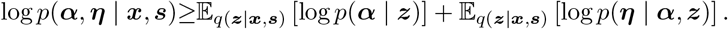

By the conditional independence assumption in Eq. 4:

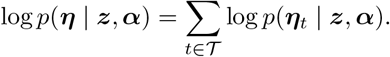

#### Modeling of *q* (***z*** | ***x, s***)

The variational posterior is chosen to be Gaussian with a diagonal covariance matrix, with parameters given by a Multi-Layer Perceptron (MLP) encoder applied to (***x, s***).

#### Training procedure

We use the analytical expressions for the Kullback-Leibler divergence and the log-likelihoods to optimize the lower bound over generative and variational parameters.

The standard VAE terms are

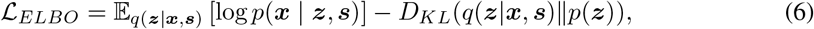

while the spatial contribution is

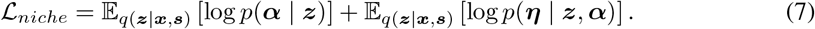

Finally, we have added a simple linear classifier that predicts the cell type from the latent space and was trained by cross-entropy minimization. Introduced in scANVI (Xu et al., 2021), it proved helpful in increasing separation between different cell types in the latent space.

Introducing *ρ >* 0 as the classification ratio and *β >* 0 as the spatial regularization strength, we write the total loss as

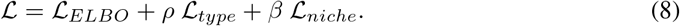

### B Implementation details

In practice, the niche cell type composition and average expression are computed using the *K* nearest neighbors of each cell; in our experiments, we fixed *K* = 20. We use scANVI (Xu et al., 2021) to learn expression embeddings ***η***. The niche composition decoder and the niche expression decoder have 1 hidden layer of size 128. During training, the learning rate is 5 *×* 10^−4^ and we use Adam as optimizer (Kingma & Ba, 2014). We warm up the KL weight from 0 to 1 for 400 epochs. Regarding the loss weights defined in Eq. 8, the spatial regularization strength is set to *β* = 10, as it led to the best tradeoff between cell type preservation and integration of niche information, while *ρ* = 20. We trained for 1000 epochs with early stopping and a batch size of 1024 cells (brain atlas, breast cancer datasets) or 4096 cells (pancancer data).

### C Benchmarks

#### scANVI

The model has 1 hidden layer with 128 neurons and a 10-dimensional latent space. The learning rate is 1 *×* 10^−4^ and we use Adam as optimizer. We warm up the KL weight from 0 to 1 for 400 epochs. It is trained for 1000 epochs with early stopping and a batch size of 512 cells (brain atlas, breast cancer datasets) or 4096 cells (pancancer data). We use a Poisson likelihood.

#### simVI

The spatial graph is also built using a *K*-nn scheme with 20 neighbors. We kept default settings for the architecture: two hidden layers for the gene expression encoder and one for the decoder, one graph attention layer for the spatial encoder. Hidden layer size is 128. During training, mutual information regularization strength is kept to its default value of 5 and the two KL terms are kept to 1.

#### BANKSY

We run BANKSY using the default parameters described in Singhal et al. (2024), namely using a Gaussian decay as spatial weights and the Gabor filter as niche feature. The number of principal components for the dimensionality reduction is kept to 20. We ran the algorithm for *λ* = 0.2, the value that the authors chose for “spatially aware cell-typing”.

#### Nicheformer

We follow the instructions in github.com/theislab/nicheformer/get embeddings.ipynb to extract embeddings using the pre-trained model weights. For fine-tuning, we follow the relevant script github.com/theislab/nicheformer/pretraining fine tune.py and train for 5 epochs with a batch size of 12, a chunk size of 100 cells, a learning rate of 10^−5^ and a warmup of 1 step.

### D Hotspot analysis

We use Hotspot (DeTomaso & Yosef, 2021) to uncover spatial signatures of cell states. To generate Figure 3, we ran Hotspot using scVIVA latent as similarity metric, following hotspot.readthedocs.io/tutorial. We displayed the spatial expression of example genes in the thalamus, hypothalamus, and striatum modules. We processed the counts as follows: we applied a `scanpy.pp.log1p` transformation followed by min-max scaling, then selected the 99th percentile as the upper limit of the color scale.

Then, we compared the spatial patterns of gene modules obtained with scVIVA and scANVI. For each method, we first standardized the cells × modules matrix of module *×* scores, then ran Hotspot on the spatial coordinates, setting `model=`“`none`”. It outputs an autocorrelation score for each module. We visually group modules corresponding to the same spatial locations (Figures S3 and S4)) and compute a Wilcoxon signed-rank test between the two distributions of modules spatial autocorrelations, using `scipy.stats.wilcoxon` with `alternative=`“`greater`” for scVIVA scores. We report the *p*-value. Considering the same subset of matched modules, we also tested how much the module scores predicted the main region label: we computed the ROC curves between module scores and region labels, then the ROC area scores (AUC), outputting a modules *×* regions AUC matrix. We then reported the maximum AUC for each module, leading to distributions of module AUC for scANVI and scVIVA. We compared these distributions using the same Wilcoxon test and alternative hypothesis.

To generate Figure 4B-C, we applied Hotspot using the same parameters as for Figure 3, but using the spatial coordinates. Then we used the module scores to assign cells to modules by considering the maximal module score for the cell. To account for cells expressing a mixture of gene modules with no clear over-expression of one module, we set a threshold *τ* to the score. Formally, for *N* cells and *M* modules, we denote ***H*** *∈* ℝ^N ×M^ the module scores matrix. For any cell *n*, the module *m*_*n*_ is:

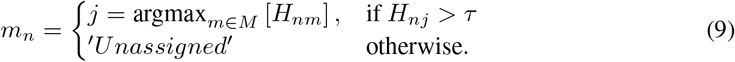

We set *τ* = 0.5. Spatial plots of module assignments *m* are displayed in Figure S5.

### E Differential Expression

#### 2-step differential expression method

In this paragraph, we explain how we leverage the generative part of scVIVA to perform differential expression (DE). Let *C1, C2* be any two groups of cells. They can correspond to different types or to the same cell type in two different spatial contexts (for instance, astrocytes in two brain regions). Our goal is to determine which genes have different expression levels between the two groups. The naive approach consists of comparing *C1* and *C2* only. However, due to errors in molecule assignment to cells, this approach returns spurious discoveries, leading to the detection of genes expressed by neighboring cells. Therefore, we should also consider the neighborhood gene expression.

To formalize this, let us consider a tissue slide *b* consisting of *N*_*b*_ cells. We compute the slide adjacency matrix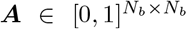, by defining either a radius neighbors graph or a nearest neighbors graph. Then, we multiply this matrix with a cell type mask to get an adjusted matrix 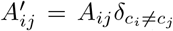, with *c*_*i*_, *c*_*j*_ being the types of cells *i* and *j*, respectively. If the dataset consists of multiple slides, we construct a block diagonal adjacency matrix containing all cells *N*. We then extract the rows corresponding to the group of interest and gather all the non-zero indices appearing in these rows. We call this list *I*_*1*_ for *C1*. Finally, we define *N1* = {*i* | *i ∈ I*_*1*_} as the set of unique indices appearing in *I*_*1*_.

To determine the upregulated genes of *C1* versus *C2*, we compute DE between {*C1, C2*}, {*N1, C2*} and {*C1, N1*}. scVIVA inherits from scvi-tools built-in differential expression module, lvm-DE (Boyeau et al., 2023), which leverages the fitted generative model to estimate log-fold changes between conditions from the normalized expression distribution. For each comparison, the Log Fold-Change (LFC) is computed as in Equation 8 of Boyeau et al. (2023), and the probability *p*_*g*_ that a gene *g* is differentially expressed is computed as in Equation 10.

The significantly upregulated genes for {*C1, N1*} define a set of local cell type markers, denoted 𝒮_1_. Conversely, if a gene is both higher expressed in *N1* compared to *C1* and *C1* compared to *C2*, it is likely that the increased expression in *C1* is spurious.

We argue that the probability of a gene being a *local marker* could be a relevant score to filter spurious genes. To compute this score, we considered the upregulation of a gene in one group relative to the upregulation in its neighborhood: a local marker *g* should verify

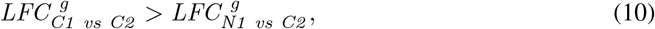

which means that the signal comes from cells in *C1* rather than their neighbors *N1*. We select genes for which *LFC*_*C1 vs C2*_ *>* 0 and use the genes 𝒮_1_ as truly differentially expressed. We also define 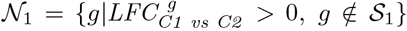. We train a Gaussian process classifier on ***X*** = [*LFC*_*C*1 *vs C*2_, *LFC*_*N*1 *vs G*2_] to classify between the *local markers* 𝒮_1_ and the *neighborhood genes N*_1_. Once fitted, the classifier returns a local marker probability 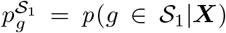 for each gene *g*, that we can compare to a given threshold Δ to filter the neighborhood genes. Algorithm 1 summarizes the procedure.

To determine the upregulated genes of *C2 vs C1*, we follow the same steps: we first retrieve *N2*, then compute DE between {*C2, C1*}, {*N2, C1*} and {*C2, N2*}.

#### Implementation details

We used the Scikit-learn (Pedregosa et al., 2011) Gaussian process classifier implementation. We defined the kernel as the product of a constant kernel *C* with a rational quadratic kernel 𝒦 (*l, a*). We tuned the Gaussian process hyperparameters by sampling 20 combinations within given bounds. We set *C ∈* [10^−3^, 10^3^], *l ∈* [10, 10^2^] and *a ∈* [10^−3^, 1].

We first applied this method to endothelial cells in breast cancer (Figures 4 and S6), with *C1, C2* obtained using maximum Hotspot module scores as detailed in Section D. We ran lvm-DE with the following parameters: a pseudo-count *ϵ* = 10^−4^, a LFC cutoff *δ* = 0.03 except for the {*C1, N1*} comparison where *δ* = 0.15, *n*_*samples*_ = 10^5^ samples from the posterior and a FDR of 0.2. We defined neighborhoods by computing the nearest neighbors graph with *k*_*DE*_ = 6 neighbors, which led to an average number of neighbors *k*_*adj*_ = 2.7 after removing cell-type connections. In the Figure 4E (resp. S6C), we plotted the set of genes 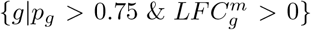 for the {*C1 vs C2*} (resp. {*C2 vs C1*}) comparisons, where 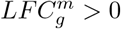 denotes the median log fold change across the *n*_*samples*_. These genes are colored by their local marker/ neighborhood gene label and we also display the marker gene probabilities 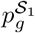.

Then, we wanted to quantify the reproducibility of the local markers results between the two different cell segmentations and the number of neighborhood genes discarded after the re-segmentation. For this, we computed Jaccard indices between sets of local markers and neighborhood genes.

For the spatial plots of gene expression, we processed the counts as follows: we applied a `scanpy.pp.log1p` transformation followed by min-max scaling, then selected the 99.9th percentile as the upper limit of the color scale.

To generate Figure 5, we first obtained the *C1, C2* groups by clustering the scVIVA latent space, either restricted to CD8+ T cells or Tregs, using the k-means algorithm (k=2). We ran lvm-DE with the same parameters as in Figures 4 except a pseudo-count *ϵ* = 10^−2^. In Figure 5F, we plotted the set of genes 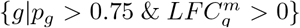, where 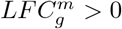 denotes the median log fold change across the *n*_*samples*_. In Figure 5H, we plotted the set of genes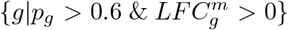. These genes are colored by their local marker/ neighborhood gene label, and we also display the marker gene probabilities 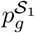. We used the same transformations as in Figures 4 and S6 for the spatial plots of expression.

##### Algorithm 1 Find robust upregulated genes of *C1* vs *C2* using the 2-step DE

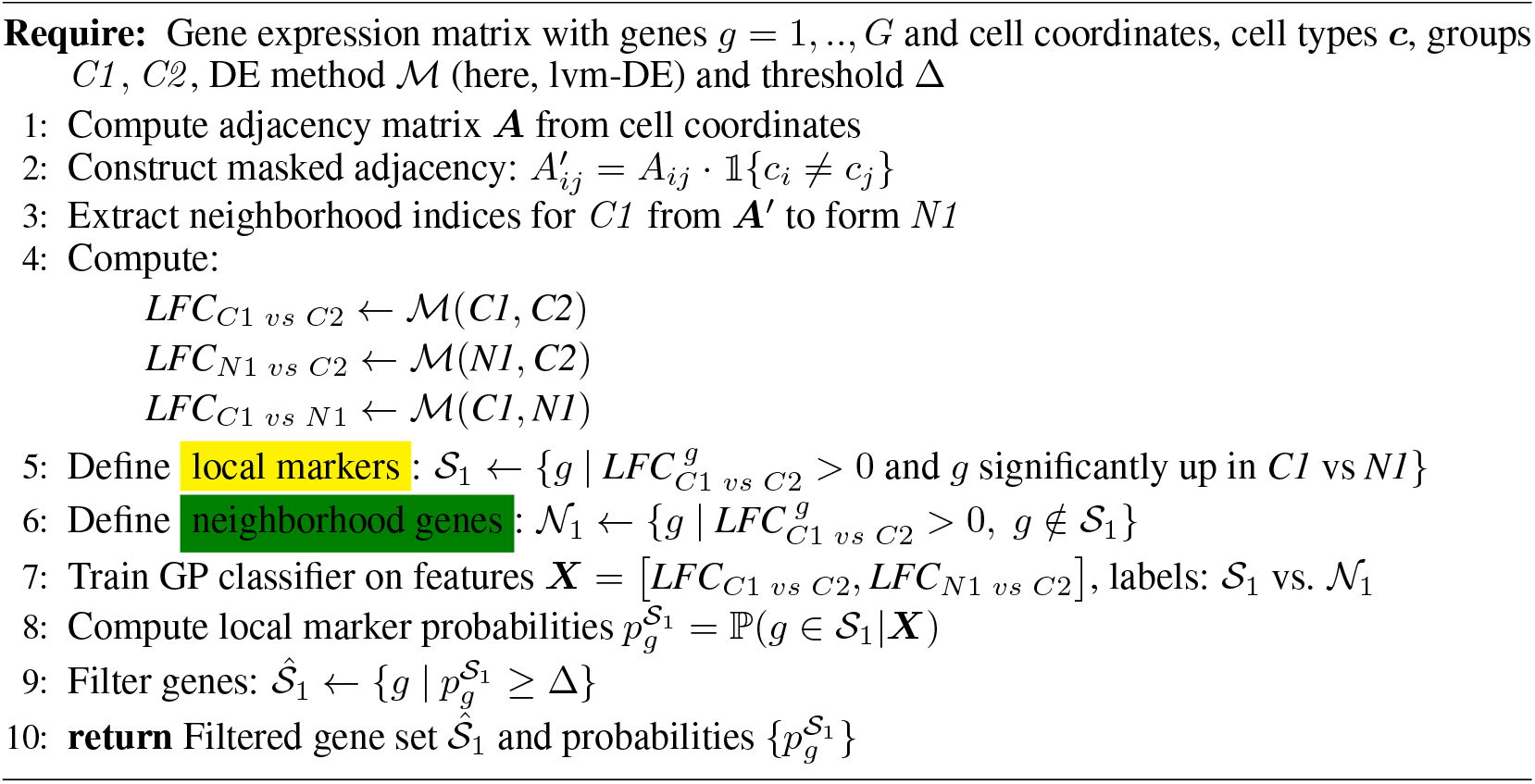

### F Datasets

All data sets used in this study are publicly available. We provide download links and processing steps below.

#### MERFISH mouse brain atlas

Zhang et al. (2023) imaged a panel of 1100 genes across the entire adult mouse brain with the MERFISH technology. For two different mice, we selected three consecutive slices, gathering six slices and 250k cells. We used the provided cell type and major brain region annotations. The data can be downloaded from the CellXGene portal. Our subset corresponds to slide indices ^*′*^*C*57*BL*6*J* − 1.074^*′*^,^*′*^ *C*57*BL*6*J* − 1.077^*′*^,^*′*^ *C*57*BL*6*J* − 1.079^*′*^,^*′*^ *C*57*BL*6*J* − 2.035^*′*^,^*′*^ *C*57*BL*6*J* − 2.036^*′*^,^*′*^ *C*57*BL*6*J* − 2.037^*′*^.

#### XENIUM human breast cancer

Using the 10X XENIUM In Situ platform in human breast cancer samples, Janesick et al. (2023) studied tumor invasion in ductal carcinoma in situ (DCIS). Two tumor samples were analyzed with a panel of 313 genes for 282k cells. The data can be downloaded from 10x Genomics. We re-segmented the cells using Proseg (Jones et al., 2024), which resulted in 279k cells.

#### Vizgen pan-cancer atlas

We assembled the 8 different cancer types of the MERSCOPE FFPE immuno-oncology release, downloadable from Vizgen. We re-segmented the cells using Proseg, leading to 10 373 000 cells *×* 550 genes. To assign coarse cell types, we performed Louvain clustering on a random subset of 50% of the cells, followed by manual annotation of the clusters.

These annotations were then propagated to the full dataset using resolVI (Ergen & Yosef, 2025) with default parameters. As our analysis focuses on T cells, we further refined the annotations by removing cells with fewer than 50 counts and subclustering the scVIVA T cell embedding. We applied Leiden clustering at a resolution of 0.5, identifying 12 distinct clusters (FigureS7A). We then performed one-vs-all differential expression analysis using the Wilcoxon test as implemented in `scanpy.rank_gene_groups` (Figure S7B). Finally, we filtered out T cells with mixed phenotypes, designating them as low-quality, and restricted the remaining analysis to high-quality T cells.

## Code availability

scVIVA is available within scvi-tools. Code to reproduce the figures can be found at github.com/LevyNat/scVIVA-reproducibility.

## Acknowledgments

We thank Martin Kim, Ori Kronfeld, Allon Wagner, Linor Bengal and Oier Etxezarreta Arrastoa for valuable discussions and feedback on this project. We also thank Shai Bagon and Romain Lopez for their advice on model design. This research was partially supported by the Israeli Council for Higher Education (CHE) via the Weizmann Data Science Research Center. The research of B.N. was partially supported by ISF grant 2362/22. Figure 1 is created with BioRender.

## 4 Supplementary figures

**Figure S1.**
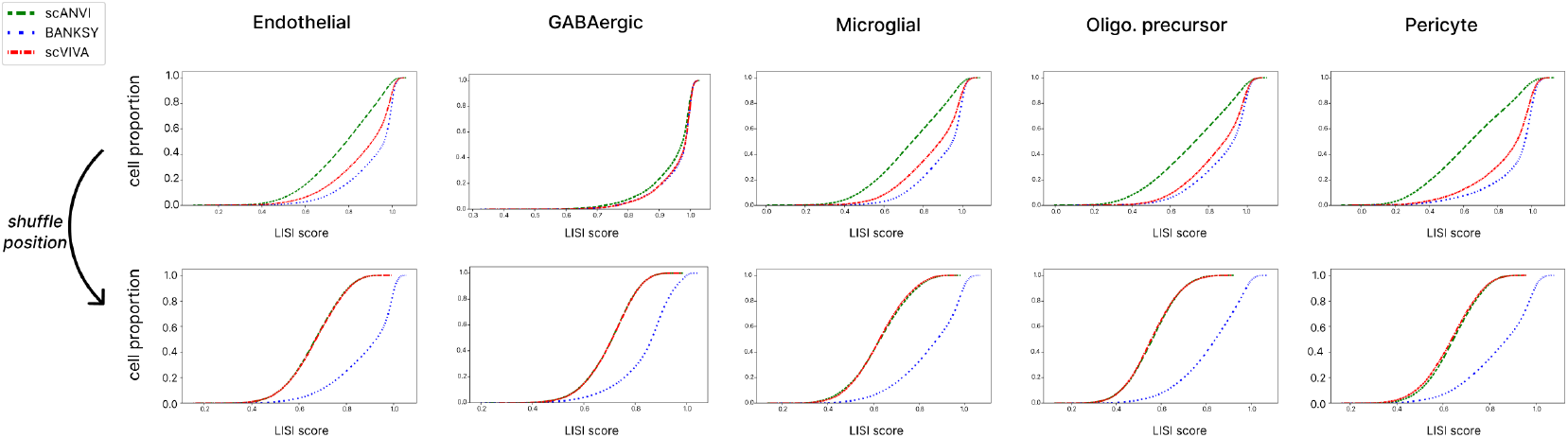
Cell type spatial coordinates shuffling experiment in the mouse brain dataset. We restrict the computation to cell types with at least 400 occurrences in 2 different regions. Distribution of cell LISI scores for each cell type before and after shuffling.

**Figure S2.**
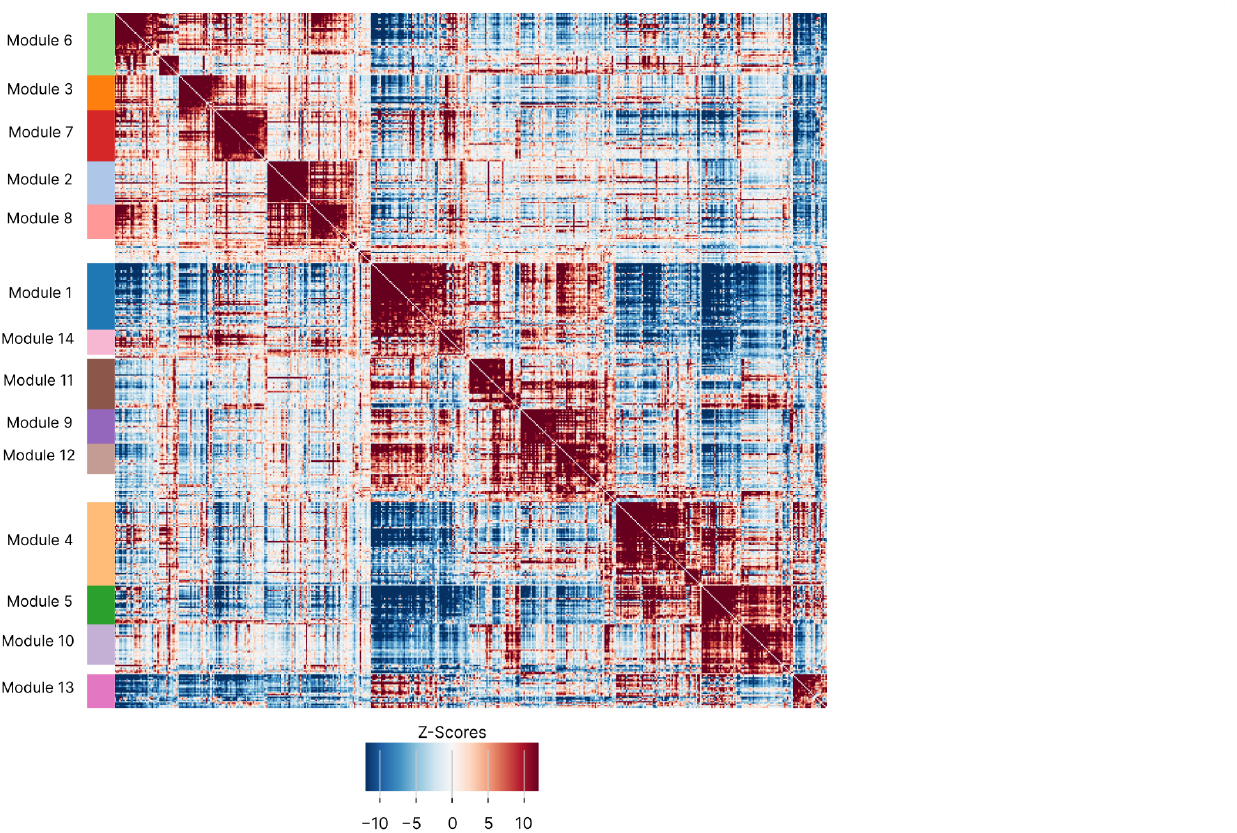
Hotspot analysis of the mouse brain dataset. Local gene-gene correlations grouped by modules, computed with Hotspot applied to scVIVA latent.

**Figure S3.**
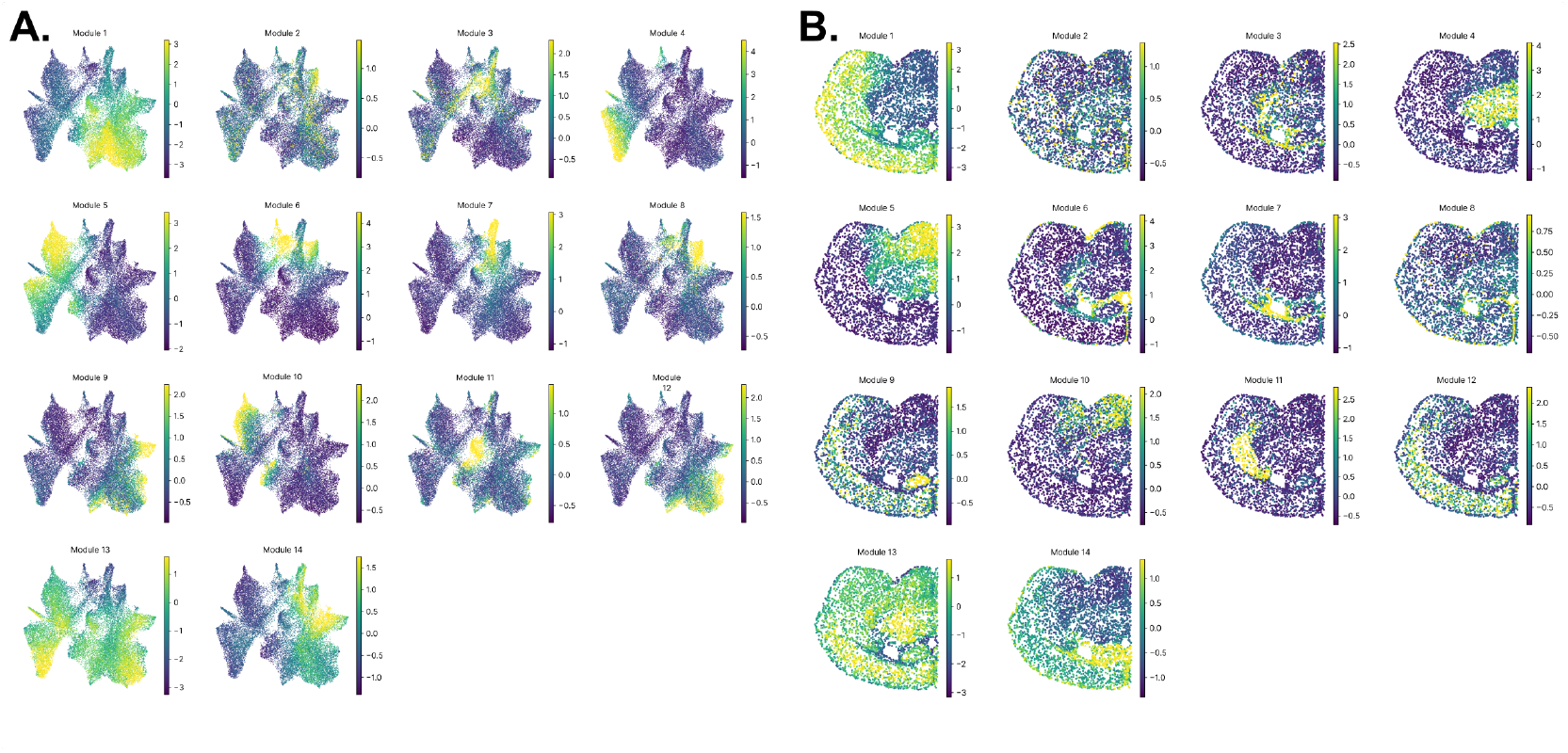
Hotspot analysis of the mouse brain dataset. Gene modules uncovered with Hotspot applied to scVIVA latent, in UMAP space **(A)** and in spatial coordinates **(B)**.

**Figure S4.**
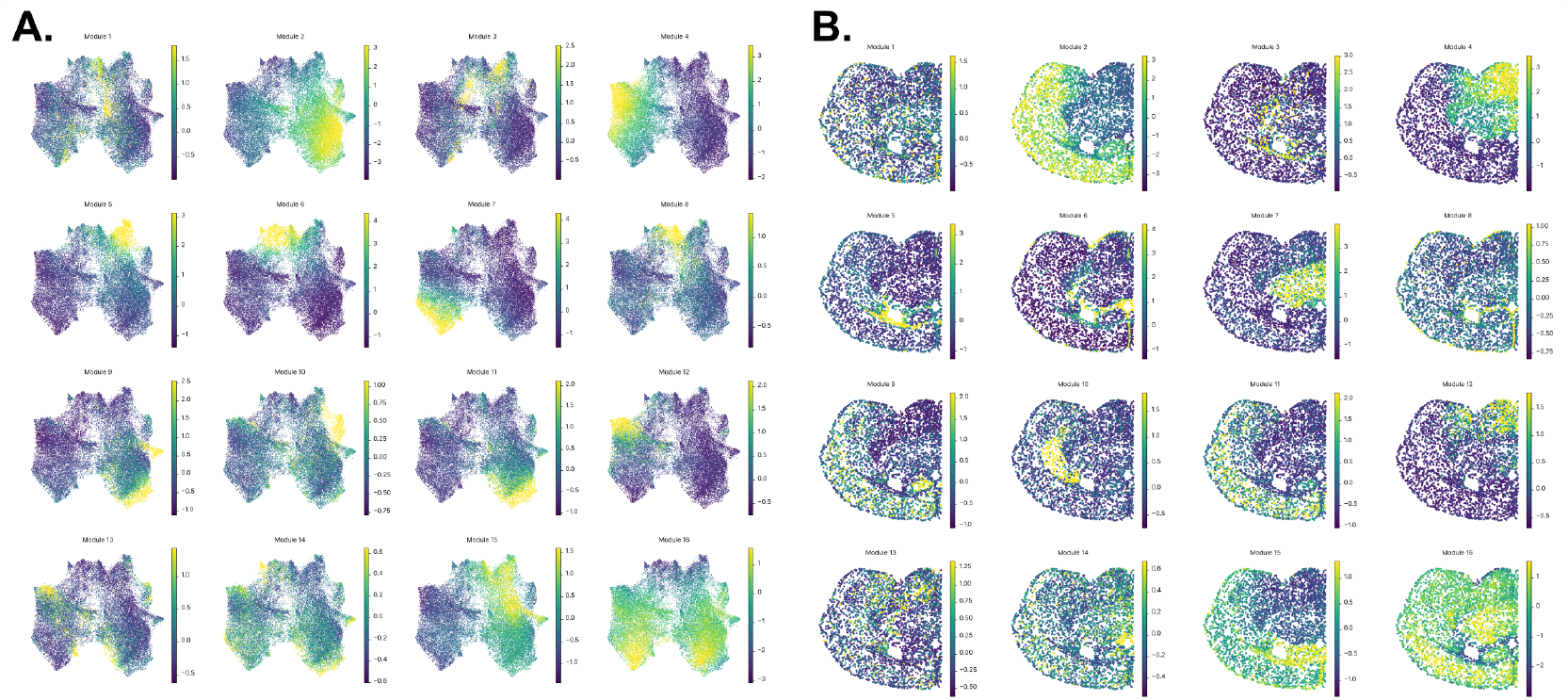
Hotspot analysis of the mouse brain dataset. Gene modules uncovered with Hotspot applied to scANVI latent, in UMAP space **(A)** and in spatial coordinates **(B)**.

**Figure S5.**
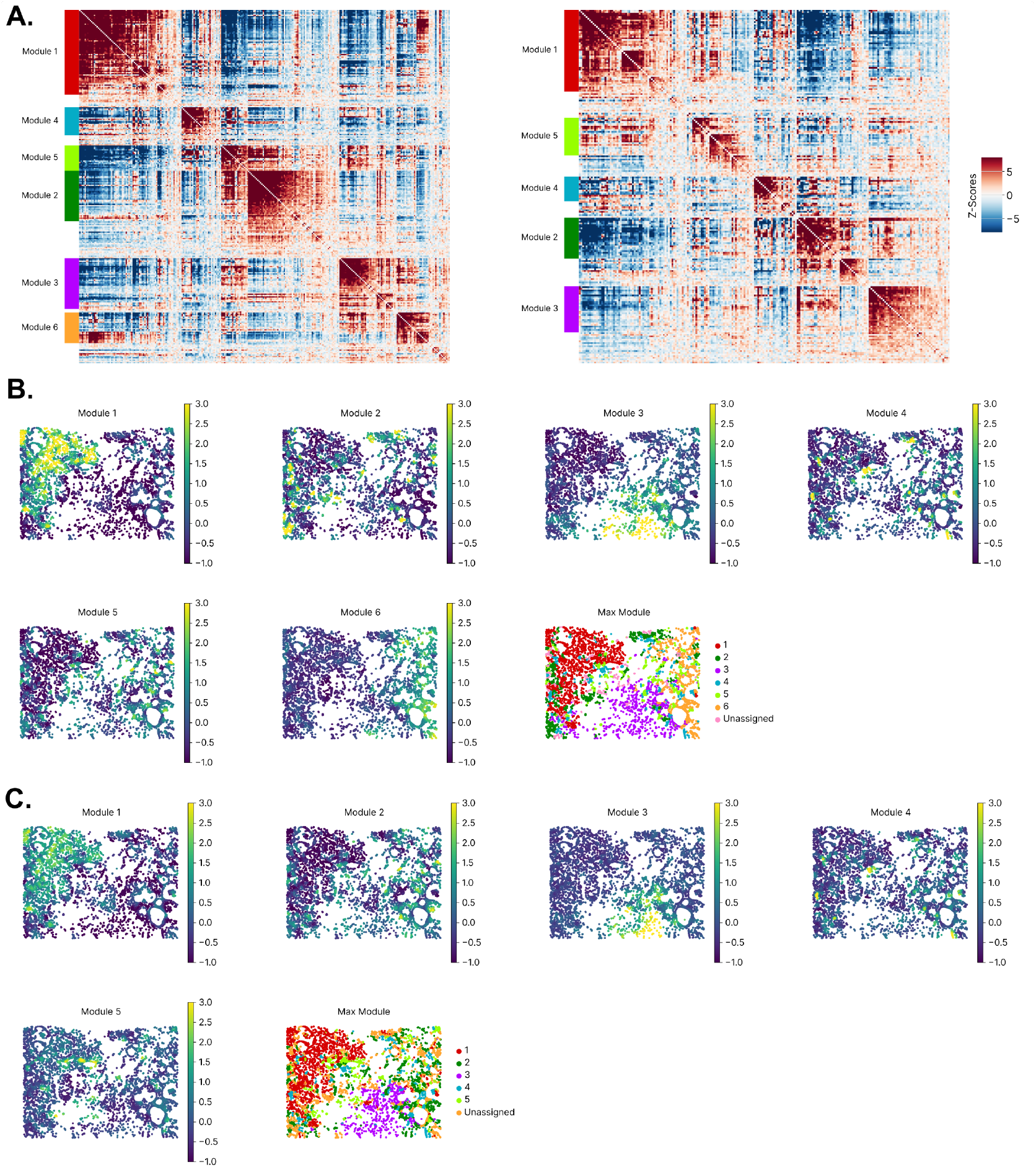
Hotspot analysis of the breast cancer dataset. **A**. Local gene-gene correlations grouped by modules, computed with Hotspot applied to the spatial coordinates, using the original segmentation (left) and after re-segmenting the cells using Proseg (right). **B**. Spatial plots of module scores. We assigned each cell to a specific module by taking the argmax over the cell module scores and setting a threshold of 0.5, under which cells are assigned to a ‘weak’ module class. **C**. Same as **B**, for the re-segmented data.

**Figure S6.**
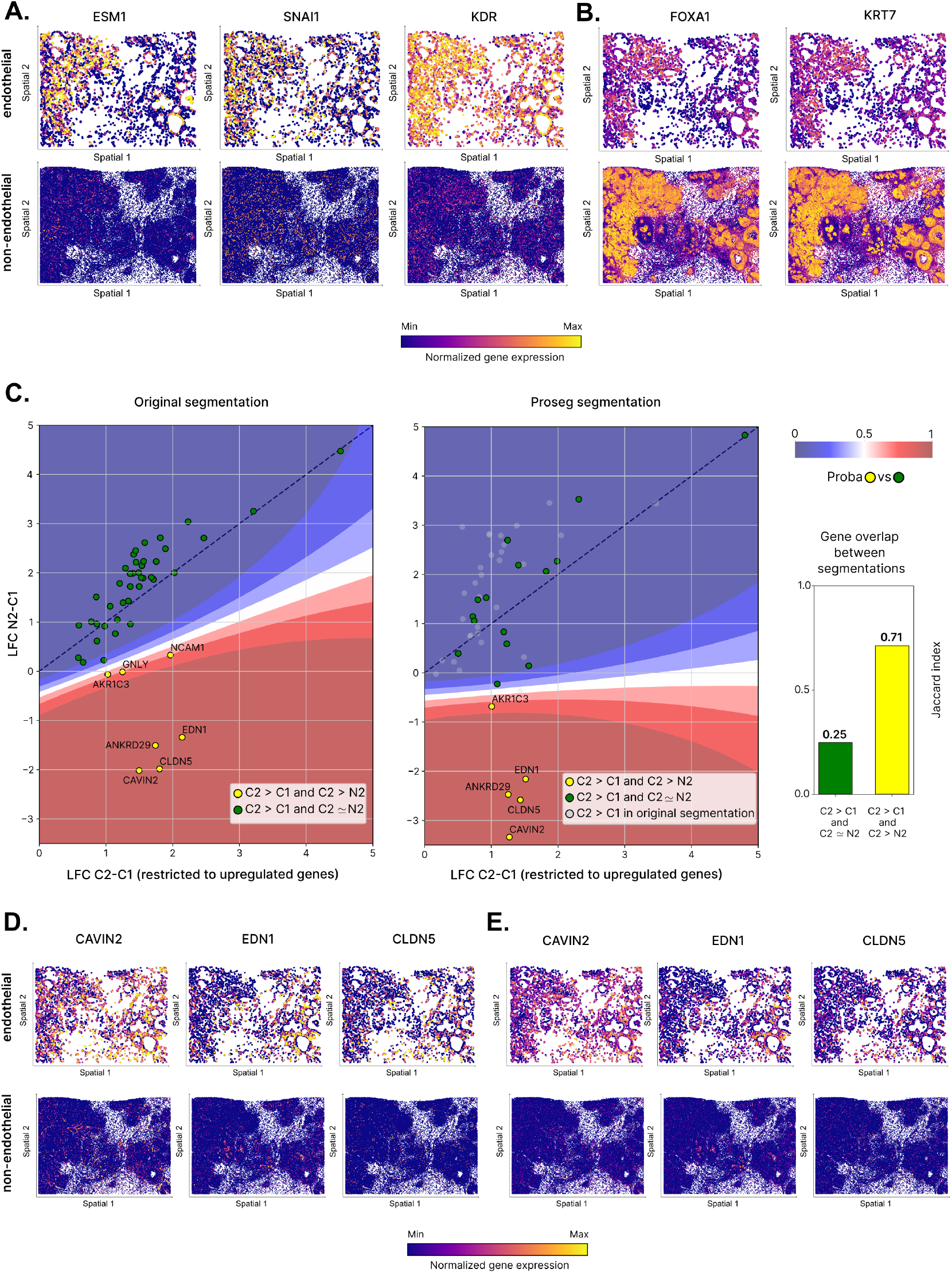
**A**. Spatial gene expression of marker genes upregulated in the invasive niche, subset to endothelial cells (top) and subset to non-endothelial cells (bottom). These genes are higher expressed in endothelial cells. Here, we display the observed expression in the original segmentation. **B**. Similar to A. These genes are higher expressed in non-endothelial cells. **C**. Median Log-Fold Change (LFC) of upregulated genes in *C2* vs *C1*, using the original segmentation (left) and the re-segmented data (right) displayed on the x-axis, while we compare differential expression computed between *N2* and *C1* on the y-axis. Genes are colored by their marker label. The classifier decision boundary is shown, and the dashed line represents the identity line. Gene overlaps for the sets of markers/ neighborhood genes are measured by Jaccard index. **D**. Similar to A., for genes overexpressed in the stroma. **E**. Similar to D., with the Proseg segmentation.

**Figure S7.**
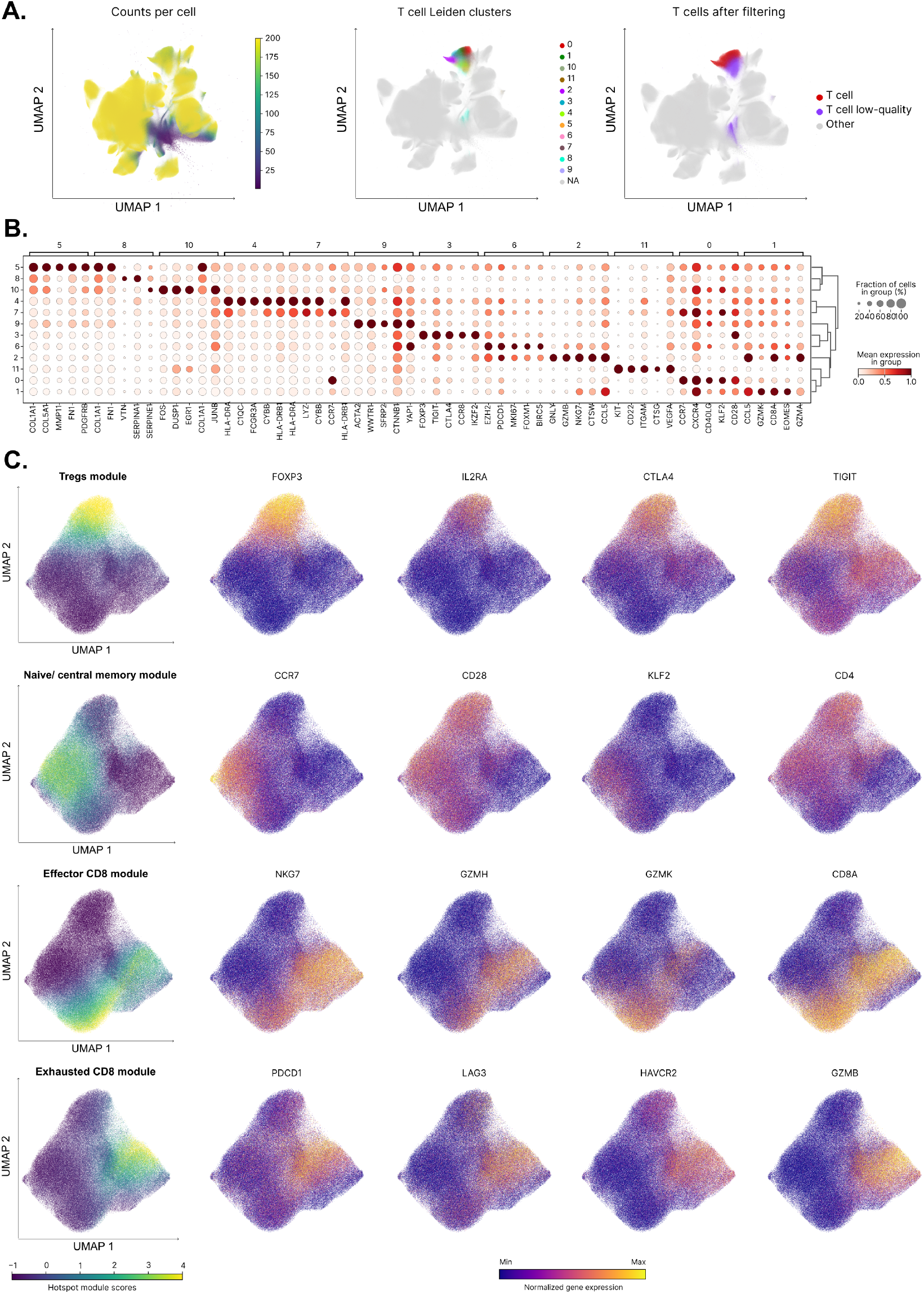
**A**. Filtering low-quality T cells. We first exclude cells with fewer than 50 counts (left) and then apply Leiden clustering on the scVIVA latent space (right). In **B**, we identify the top five marker genes for each cluster and remove clusters where non-T-cell markers dominate. This results in a set of high-confidence T cells (right). **B**. One-vs-all differential expression analysis of the Leiden clusters from **A**. As several clusters show signs of potential contamination, we retain only clusters “0”, “1”, “2”, “3”, and “6”, where the top five markers correspond to known T-cell subtypes. **C**. We run Hotspot on scVIVA embedding restricted to high-confidence T cells. Four gene modules correspond to known T cell subtypes (Figure5B). On a UMAP representation of the scVIVA embedding, we visualize the four module scores and the expression of selected top module genes.

## References

1. C. Aitken, V. Mehta, M. A. Schwartz, and E. Tzima. Mechanisms of endothelial flow sensing. Nature Cardiovascular Research, 2(6):517–529, Jun 2023. doi: 10.1038/s44161-023-00276-0.

2. Sarah Amato, Monica Averna, Diego Guidolin, Cristina Ceccoli, Elena Gatta, Simona Candiani, Marco Pedrazzi, Michela Capraro, Guido Maura, Luigi F Agnati, et al. Heteromerization of dopamine d2 and oxytocin receptor in adult striatal astrocytes. International Journal of Molecular Sciences, 24(5):4677, 2023.

3. Aurelio Balsalobre and Jacques Drouin. Pioneer factors as master regulators of the epigenome and cell fate. Nat. Rev. Mol. Cell Biol., 23(7):449–464, July 2022.

4. Poornima Bhat-Nakshatri, Hongyu Gao, Aditi S Khatpe, Adedeji K Adebayo, Patrick C McGuire, Cihat Erdogan, Duojiao Chen, Guanglong Jiang, Felicia New, Rana German, Lydia Emmert, George Sandusky, Anna Maria Storniolo, Yunlong Liu, and Harikrishna Nakshatri. Single-nucleus chromatin accessibility and transcriptomic map of breast tissues of women of diverse genetic ancestry. Nat. Med., 30(12):3482–3494, December 2024.

5. G. T. K. Boopathy, M. Kulkarni, S. Y. Ho, A. Boey, E. W. M. Chua, V. A. Barathi, T. J. Carney, X. Wang, and W. Hong. Cavin-2 regulates the activity and stability of endothelial nitric-oxide synthase (enos) in angiogenesis. Journal of Biological Chemistry, 292(43):17760–17776, Oct 2017. doi: 10.1074/jbc.M117.794743.

6. Pierre Boyeau, Jeffrey Regier, Adam Gayoso, Michael I Jordan, Romain Lopez, and Nir Yosef. An empirical bayes method for differential expression analysis of single cells with deep generative models. Proceedings of the National Academy of Sciences, 120(21):e2209124120, 2023.

7. Pierre Boyeau, Justin Hong, Adam Gayoso, Martin Kim, José L McFaline-Figueroa, Michael I Jordan, Elham Azizi, Can Ergen, and Nir Yosef. Deep generative modeling of sample-level heterogeneity in single-cell genomics. BioRxiv, pp. 2022–10, 2024.

8. Shannon K Bromley, Seddon Y Thomas, and Andrew D Luster. Chemokine receptor ccr7 guides t cell exit from peripheral tissues and entry into afferent lymphatics. Nature immunology, 6(9): 895–901, 2005.

9. Michael D Buck, Ryan T Sowell, Susan M Kaech, and Erika L Pearce. Metabolic instruction of immunity. Cell, 169(4):570–586, 2017.

10. Marcus Buggert, David A Price, Laura K Mackay, and Michael R Betts. Human circulating and tissue-resident memory CD8+ T cells. Nat. Immunol., 24(7):1076–1086, July 2023.

11. David Cabrerizo-Granados, Raúl Peña, Laura Palacios, Laura Carrillo-Bosch, Josep Lloreta-Trull, Laura Comerma, Mar Iglesias, and Antonio García de Herreros. Snail1 expression in endothelial cells controls growth, angiogenesis and differentiation of breast tumors. Theranostics, 11(16):7671, 2021.

12. Paolo Cadinu, Kisha N Sivanathan, Aditya Misra, Rosalind J Xu, Davide Mangani, Evan Yang, Joseph M Rone, Katherine Tooley, Yoon-Chul Kye, Lloyd Bod, et al. Charting the cellular biogeography in colitis reveals fibroblast trajectories and coordinated spatial remodeling. Cell, 187(8):2010–2028, 2024.

13. Anna K Casasent, Mathilde M Almekinders, Charlotta Mulder, Proteeti Bhattacharjee, Deborah Collyar, Alastair M Thompson, Jos Jonkers, Esther H Lips, Jacco van Rheenen, E Shelley Hwang, Serena Nik-Zainal, Nicholas E Navin, Jelle Wesseling, and Grand Challenge PRECISION Consortium. Learning to distinguish progressive and non-progressive ductal carcinoma in situ. Nat. Rev. Cancer, 22(12):663–678, December 2022.

14. Qian Chen, Meiying Shen, Min Yan, Xiaojian Han, Song Mu, Ya Li, Luo Li, Yingming Wang, Shenglong Li, Tingting Li, et al. Targeting tumor-infiltrating ccr8+ regulatory t cells induces antitumor immunity through functional restoration of cd4+ tconvs and cd8+ t cells in colorectal cancer. Journal of Translational Medicine, 22(1):709, 2024.

15. Anthony R Cillo, Carly Cardello, Feng Shan, Lilit Karapetyan, Sheryl Kunning, Cindy Sander, Elizabeth Rush, Arivarasan Karunamurthy, Ryan C Massa, Anjali Rohatgi, Creg J Workman, John M Kirkwood, Tullia C Bruno, and Dario AA Vignali. Blockade of LAG-3 and PD-1 leads to co-expression of cytotoxic and exhaustion gene modules in CD8+ T cells to promote antitumor immunity. Cell, 187(16):4373–4388.e15, August 2024.

16. David DeTomaso and Nir Yosef. Hotspot identifies informative gene modules across modalities of single-cell genomics. Cell systems, 12(5):446–456, 2021.

17. Mingze Dong, Harriet Kluger, Rong Fan, and Yuval Kluger. Simvi reveals intrinsic and spatial-induced states in spatial omics data. bioRxiv, 2023.

18. Rasa Elmentaite, Cecilia Domínguez Conde, Lu Yang, and Sarah A Teichmann. Single-cell atlases: shared and tissue-specific cell types across human organs. Nat. Rev. Genet., 23(7):395–410, July 2022.

19. Fumito Endo, Atsushi Kasai, Joselyn S Soto, Xinzhu Yu, Zhe Qu, Hitoshi Hashimoto, Viviana Gradinaru, Riki Kawaguchi, and Baljit S Khakh. Molecular basis of astrocyte diversity and morphology across the cns in health and disease. Science, 378(6619):eadc9020, 2022.

20. Can Ergen and Nir Yosef. Resolvi-addressing noise and bias in spatial transcriptomics. bioRxiv, pp. 2025–01, 2025.

21. Jéròme Galon and Daniela Bruni. Approaches to treat immune hot, altered and cold tumours with combination immunotherapies. Nature reviews Drug discovery, 18(3):197–218, 2019.

22. V. Geldhof, L. P. M. H. de Rooij, L. Sokol, J. Amersfoort, M. De Schepper, K. Rohlenova, G. Hoste, A. Vanderstichele, A. M. Delsupehe, E. Isnaldi, N. Dai, F. Taverna, S. Khan, A. K. Truong, L. A. Teuwen, F. Richard, L. Treps, A. Smeets, I. Nevelsteen, B. Weynand, S. Vinckier, L. Schoonjans, J. Kalucka, C. Desmedt, P. Neven, M. Mazzone, G. Floris, K. Punie, M. Dewerchin, G. Eelen, H. Wildiers, X. Li, Y. Luo, and P. Carmeliet. Single cell atlas identifies lipid-processing and immunomodulatory endothelial cells in healthy and malignant breast. Nature Communications, 13(1):5511, sep 2022. doi: 10.1038/s41467-022-33052-y.

23. Michael A Harris, Peter Savas, Balaji Virassamy, Megan MR O’Malley, Jasmine Kay, Scott N Mueller, Laura K Mackay, Roberto Salgado, and Sherene Loi. Towards targeting the breast cancer immune microenvironment. Nat. Rev. Cancer, 24(8):554–577, August 2024.

24. Doron Haviv, Ján Remšík, Mohamed Gatie, Catherine Snopkowski, Meril Takizawa, Nathan Pereira, John Bashkin, Stevan Jovanovich, Tal Nawy, Ronan Chaligne, et al. The covariance environment defines cellular niches for spatial inference. Nature Biotechnology, pp. 1–12, 2024.

25. Á. Herrero-Navarro, L. Puche-Aroca, V. Moreno-Juan, A. Sempere-Ferràndez, A. Espinosa, R. Susín, L. Torres-Masjoan, E. Leyva-Díaz, M. Karow, M. Figueres-Oñate, L. López-Mascaraque, J.P. López-Atalaya, B. Berninger, and G. López-Bendito. Astrocytes and neurons share region-specific transcriptional signatures that confer regional identity to neuronal reprogramming. Science Advances, 7(15):eabe8978, 2021.

26. Helene Hoffmann, Martin Wartenberg, Sandra Vorlova, Franziska Karl-Schöller, Matthias Kallius, Oliver Reinhardt, Asli Öztürk, Leah S Schuhmair, Verena Burkhardt, Sabine Gôtzner, et al. Normalization of snai1-mediated vessel dysfunction increases drug response in cancer. Oncogene, 43(35):2661–2676, 2024.

27. Amanda Janesick, Robert Shelansky, Andrew D Gottscho, Florian Wagner, Stephen R Williams, Morgane Rouault, Ghezal Beliakoff, Carolyn A Morrison, Michelli F Oliveira, Jordan T Sicherman, et al. High resolution mapping of the tumor microenvironment using integrated single-cell, spatial and in situ analysis. Nature Communications, 14(1):8353, 2023.

28. Nicole Joller, Ester Lozano, Patrick R Burkett, Bonny Patel, Sheng Xiao, Chen Zhu, Junrong Xia, Tze G Tan, Esen Sefik, Vijay Yajnik, et al. Treg cells expressing the coinhibitory molecule tigit selectively inhibit proinflammatory th1 and th17 cell responses. Immunity, 40(4):569–581, 2014.

29. Daniel C Jones, Anna E Elz, Azadeh Hadadianpour, Heeju Ryu, David R Glass, and Evan W Newell. Cell simulation as cell segmentation. bioRxiv, 2024.

30. Sjoerd D Joustra, Onno C Meijer, Charlotte A Heinen, Isabel M Mol, El Houari Laghmani, Rozemarijn MA Sengers, Gabriela Carreno, AS van Trotsenburg, Nienke R Biermasz, Daniel J Bernard, et al. Spatial and temporal expression of immunoglobulin superfamily member 1 in the rat. J Endocrinol, 226(3):181–191, 2015.

31. Kelly Kersten, Kenneth H Hu, Alexis J Combes, Bushra Samad, Tory Harwin, Arja Ray, Arjun Arkal Rao, En Cai, Kyle Marchuk, Jordan Artichoker, et al. Spatiotemporal co-dependency between macrophages and exhausted cd8+ t cells in cancer. Cancer Cell, 40(6):624–638, 2022.

32. Baljit S Khakh. Astrocyte–neuron interactions in the striatum: insights on identity, form, and function. Trends in neurosciences, 42(9):617–630, 2019.

33. Chang N Kim, David Shin, Albert Wang, and Tomasz J Nowakowski. Spatiotemporal molecular dynamics of the developing human thalamus. Science, 382(6667):eadf9941. 2023a.

34. Rachel D Kim, Anne E Marchildon, Paul W Frazel, Philip Hasel, Amy X Guo, and Shane A Liddelow. Temporal and spatial analysis of astrocytes following stroke identifies novel drivers of reactivity. bioRxiv, 2023b.

35. Diederik P Kingma and Jimmy Ba. Adam: A method for stochastic optimization. arXiv preprint 1412.6980, 2014.

36. Diederik P Kingma and Max Welling. Auto-encoding variational bayes. arXiv preprint 1312.6114, 2013.

37. V. Kleshchevnikov, A. Shmatko, E. Dann, A. Aivazidis, H. W. King, T. Li, R. Elmentaite, A. Lomakin, V. Kedlian, A. Gayoso, M. S. Jain, J. S. Park, L. Ramona, E. Tuck, A. Arutyunyan, R. Vento-Tormo, M. Gerstung, L. James, O. Stegle, and O. A. Bayraktar. Cell2location maps fine-grained cell types in spatial transcriptomics. Nature Biotechnology, 40(5):661–671, May 2022.

38. Ilya Korsunsky, Nghia Millard, Jean Fan, Kamil Slowikowski, Fan Zhang, Kevin Wei, Yuriy Baglaenko, Michael Brenner, Po-ru Loh, and Soumya Raychaudhuri. Fast, sensitive and accurate integration of single-cell data with harmony. Nature methods, 16(12):1289–1296, 2019.

39. H. G. Lee, M. A. Wheeler, and F. J. Quintana. Function and therapeutic value of astrocytes in neurological diseases. Nature Reviews Drug Discovery, 21(5):339–358, May 2022. doi: 10.1038/s41573-022-00390-x. Epub 2022 Feb 16.

40. Chunxiao Li, Ping Jiang, Shuhua Wei, Xiaofei Xu, and Junjie Wang. Regulatory t cells in tumor microenvironment: new mechanisms, potential therapeutic strategies and future prospects. Molecular cancer, 19:1–23, 2020.

41. R. Li, J. R. Ferdinand, K. W. Loudon, G. S. Bowyer, S. Laidlaw, F. Muyas, L. Mamanova, J. B. Neves, L. Bolt, E. S. Fasouli, A. R. J. Lawson, M. D. Young, Y. Hooks, T. R. W. Oliver, T. M. Butler, J. N. Armitage, T. Aho, A. C. P. Riddick, V. Gnanapragasam, S. J. Welsh, K. B. Meyer, A. Y. Warren, M. G. B. Tran, G. D. Stewart, I. Cortés-Ciriano, S. Behjati, M. R. Clatworthy, P. J. Campbell, S. A. Teichmann, and T. J. Mitchell. Mapping single-cell transcriptomes in the intra-tumoral and associated territories of kidney cancer. Cancer Cell, 40(12):1583–1599.e10, December 2022.

42. Anastasia Litinetskaya, Maiia Shulman, Soroor Hediyeh-zadeh, Amir Ali Moinfar, Fabiola Curion, Artur Szalata, Alireza Omidi, Mohammad Lotfollahi, and Fabian J Theis. Multimodal weakly supervised learning to identify disease-specific changes in single-cell atlases. bioRxiv, pp. 2024–07, 2024.

43. Romain Lopez, Jeffrey Regier, Michael B. Cole, Michael I. Jordan, and Nir Yosef. Deep generative modeling for single-cell transcriptomics. Nature Methods, December 2018.

44. Malte D Luecken, Maren Büttner, Kridsadakorn Chaichoompu, Anna Danese, Marta Interlandi, Michaela F Müller, Daniel C Strobl, Luke Zappia, Martin Dugas, Maria Colomé-Tatché, et al. Benchmarking atlas-level data integration in single-cell genomics. Nature methods, 19(1):41–50, 2022.

45. H. Mathys, C. A. Boix, L. A. Akay, Z. Xia, J. Davila-Velderrain, A. P. Ng, X. Jiang, G. Abdelhady, K. Galani, J. Mantero, N. Band, B. T. James, S. Babu, F. Galiana-Melendez, K. Louderback, D. Prokopenko, R. E. Tanzi, D. A. Bennett, L. H. Tsai, and M. Kellis. Single-cell multiregion dissection of alzheimer’s disease. Nature, 632(8026):858–868, 2024.

46. R. J. Motzer, R. Banchereau, H. Hamidi, T. Powles, D. McDermott, M. B. Atkins, B. Escudier, L. F. Liu, N. Leng, A. R. Abbas, J. Fan, H. Koeppen, J. Lin, S. Carroll, K. Hashimoto, S. Mariathasan, M. Green, D. Tayama, P. S. Hegde, C. Schiff, M. A. Huseni, and B. Rini. Molecular subsets in renal cancer determine outcome to checkpoint and angiogenesis blockade. Cancer Cell, 38(6): 803–817.e4, December 2020. doi: 10.1016/j.ccell.2020.10.011.

47. Matthias Ollivier, Joselyn S Soto, Kay E Linker, Stefanie L Moye, Yasaman Jami-Alahmadi, Anthony E Jones, Ajit S Divakaruni, Riki Kawaguchi, James A Wohlschlegel, and Baljit S Khakh. Crym-positive striatal astrocytes gate perseverative behaviour. Nature, 627(8003):358–366, 2024.

48. Giovanni Palla, David S Fischer, Aviv Regev, and Fabian J Theis. Spatial components of molecular tissue biology. Nature Biotechnology, 40(3):308–318, 2022.

49. Fabian Pedregosa, Gaël Varoquaux, Alexandre Gramfort, Vincent Michel, Bertrand Thirion, Olivier Grisel, Mathieu Blondel, Peter Prettenhofer, Ron Weiss, Vincent Dubourg, et al. Scikit-learn: Machine learning in python. the Journal of machine Learning research, 12:2825–2830, 2011.

50. Sarah J Pfau, Urs H Langen, Theodore M Fisher, Indumathi Prakash, Faheem Nagpurwala, Ricardo A Lozoya, Wei-Chung Allen Lee, Zhuhao Wu, and Chenghua Gu. Characteristics of blood–brain barrier heterogeneity between brain regions revealed by profiling vascular and perivascular cells. Nature Neuroscience, 27(10):1892–1903, 2024.

51. L. Pérez-Gutiérrez and N. Ferrara. Biology and therapeutic targeting of vascular endothelial growth factor a. Nature Reviews Molecular Cell Biology, 24(11):816–834, November 2023. doi: 10.1038/s41580-023-00631-w.

52. A.D. Reed, S. Pensa, A. Steif, J. Stenning, D.J. Kunz, L.J. Porter, K. Hua, P. He, A.J. Twigger, A.J.Q. Siu, K. Kania, R. Barrow-McGee, I. Goulding, J.J. Gomm, V. Speirs, J.L. Jones, J.C. Marioni, and W.T. Khaled. A single-cell atlas enables mapping of homeostatic cellular shifts in the adult human breast. Nature Genetics, 56(4):652–662, Apr 2024. doi: 10.1038/s41588-024-01688-9. Epub 2024 Mar 28.

53. Vipul Singhal, Nigel Chou, Joseph Lee, and et al. Banksy unifies cell typing and tissue domain segmentation for scalable spatial omics data analysis. Nature Genetics, 2024. doi: 10.1038s41588-024-01664-3.

54. Katarzyna Sobierajska, Wojciech Michal Ciszewski, Izabela Sacewicz-Hofman, and Jolanta Niewiarowska. Endothelial cells in the tumor microenvironment. Tumor microenvironment: Non-hematopoietic cells, pp. 71–86, 2020.

55. Maria M Steele, Abhinav Jaiswal, Ines Delclaux, Ian D Dryg, Dhaarini Murugan, Julia Femel, Sunny Son, Haley du Bois, Cameron Hill, Sancy A Leachman, Young H Chang, Lisa M Coussens, Niroshana Anandasabapathy, and Amanda W Lund. T cell egress via lymphatic vessels is tuned by antigen encounter and limits tumor control. Nat. Immunol., 24(4):664–675, April 2023.

56. Lukasz Mateusz Szewczyk, Marcin Andrzej Lipiec, Ewa Liszewska, Ksenia Meyza, Joanna Urban-Ciecko, Ludwika Kondrakiewicz, Anna Goncerzewicz, Kamil Rafalko, Tomasz Grzegorz Krawczyk, Karolina Bogaj, et al. Astrocytic β-catenin signaling via tcf7l2 regulates synapse development and social behavior. Molecular Psychiatry, 29(1):57–73, 2024.

57. Vizgen. Merscope ffpe solution. URL https://info.vizgen.com/merscope-ffpe-solution. Accessed: 2025-02-27.

58. Allon Wagner, Aviv Regev, and Nir. Yosef. Revealing the vectors of cellular identity with single-cell genomics. Nature Biotechnology, 34(11):1145–1160, 2016.

59. Yuanjia Wen, Yu Xia, Xiangping Yang, Huayi Li, and Qinglei Gao. Ccr8: a promising therapeutic target against tumor-infiltrating regulatory t cells. Trends in Immunology, 2025.

60. Bo Wu, Bo Zhang, Bowen Li, Haoqi Wu, and Meixi Jiang. Cold and hot tumors: from molecular mechanisms to targeted therapy. Signal Transduction and Targeted Therapy, 9(1):274, 2024.

61. Chenling Xu, Romain Lopez, Edouard Mehlman, Jeffrey Regier, Michael I Jordan, and Nir Yosef. Probabilistic harmonization and annotation of single-cell transcriptomics data with deep generative models. Molecular systems biology, 17(1):e9620, 2021.

62. Z. Yao, C. T. J. van Velthoven, M. Kunst, M. Zhang, D. McMillen, C. Lee, W. Jung, J. Goldy, A. Abdelhak, M. Aitken, K. Baker, P. Baker, E. Barkan, D. Bertagnolli, A. Bhandiwad, C. Bielstein, P. Bishwakarma, J. Campos, D. Carey, T. Casper, A. B. Chakka, R. Chakrabarty, S. Chavan, M. Chen, M. Clark, J. Close, K. Crichton, S. Daniel, P. DiValentin, T. Dolbeare, L. Ellingwood, E. Fiabane, T. Fliss, J. Gee, J. Gerstenberger, A. Glandon, J. Gloe, J. Gould, J. Gray, N. Guilford, J. Guzman, D. Hirschstein, W. Ho, M. Hooper, M. Huang, M. Hupp, K. Jin, M. Kroll, K. Lathia, A. Leon, S. Li, B. Long, Z. Madigan, J. Malloy, J. Malone, Z. Maltzer, N. Martin, R. McCue, R. McGinty, N. Mei, J. Melchor, E. Meyerdierks, T. Mollenkopf, S. Moonsman, T. N. Nguyen, S. Otto, T. Pham, C. Rimorin, A. Ruiz, R. Sanchez, L. Sawyer, N. Shapovalova, N. Shepard, C. Slaughterbeck, J. Sulc, M. Tieu, A. Torkelson, H. Tung, N. Valera Cuevas, S. Vance, K. Wadhwani, K. Ward, B. Levi, C. Farrell, R. Young, B. Staats, M. M. Wang, C. L. Thompson, S. Mufti, C. M. Pagan, L. Kruse, N. Dee, S. M. Sunkin, L. Esposito, M. J. Hawrylycz, J. Waters, L. Ng, K. Smith, B. Tasic, X. Zhuang, and H. Zeng. A high-resolution transcriptomic and spatial atlas of cell types in the whole mouse brain. Nature, 624(7991):317–332, Dec 2023.

63. Nadav Yayon, Veronika R Kedlian, Lena Boehme, Chenqu Suo, Brianna T Wachter, Rebecca T Beuschel, Oren Amsalem, Krzysztof Polanski, Simon Koplev, Elizabeth Tuck, et al. A spatial human thymus cell atlas mapped to a continuous tissue axis. Nature, 635(8039):708–718, 2024.

64. Nir Yosef and Aviv Regev. Writ large: Genomic dissection of the effect of cellular environment on immune response. Science, 354(6308):64–68, 2016.

65. K. K. Youssef and M. A. Nieto. Epithelial-mesenchymal transition in tissue repair and degeneration. Nature Reviews Molecular Cell Biology, 25(9):720–739, September 2024. doi: 10.1038/s41580-024-00733-z.

66. Q. Zeng, M. Mousa, A. S. Nadukkandy, L. Franssens, H. Alnaqbi, F. Y. Alshamsi, H. A. Safar, and P. Carmeliet. Understanding tumour endothelial cell heterogeneity and function from single-cell omics. Nature Reviews Cancer, 23(8):544–564, August 2023. doi: 10.1038/s41568-023-00591-5.

67. Meng Zhang, Xingjie Pan, Won Jung, Aaron Halpern, Stephen W Eichhorn, Zhiyun Lei, Limor Cohen, Kimberly A Smith, Bosiljka Tasic, Zizhen Yao, et al. A molecularly defined and spatially resolved cell atlas of the whole mouse brain. Nature, pp. 343–354, 2023.

